# Constitutive immune surveillance of nasal mucosa by three neutrophil subsets with distinct origin, phenotype, and function

**DOI:** 10.1101/2024.03.06.583781

**Authors:** Rodrigo J. Gonzalez, Pavel Hanč, David Alvarez, Samuel W. Kazer, Marie-Angele Messou, Irina B. Mazo, Colette Matysiak Match, Rohit Garg, Jennifer D. Helble, Paris Pallis, Rachel Ende, Alan Basset, Rick Malley, Isabelle Derre, Michael N. Starnbach, Ulrich H. von Andrian

## Abstract

The nasal mucosa (NM) has several critical functions, including as a chemosensory organ, as a filter and conditioning surface of inhaled air for the lower airways, and as a first line of defense against airborne infections. Owing to its constant exposure to ever-changing environments, the NM is arguably the most frequently infected tissue in mammals. Consequently, vertebrates harbor an intricate network of subepithelial immune cells that are dispersed throughout the NM. However, the origin, composition, and function of nasal immune cells and their pathophysiological role are poorly understood. Here, we show that murine steady-state NM harbors a prominent population of extravascular neutrophils (EVN) that are abundant in both conventional and germ-free mice, suggesting that their presence is not driven by microbial stimuli. Nasal EVN can be subdivided into three phenotypically distinct subsets: one population that we have termed nN1 is CD11b^int^ Ly6G^int^, while the other two subsets are both CD11b^hi^ Ly6G^hi^ and distinguishable by the absence (nN2) or presence (nN3) of CD11c and SiglecF. nN1 EVN originate in bone marrow (BM) within osseous structures in the skull. These locally produced neutrophils appear to access the adjacent NM via conduits that connect BM cavities to the submucosal lamina propria. nN2 cells reach the NM via the blood and readily engulf infectious microbes. In the absence of infection, nN2 cells differentiate into the nN3 subset, which does not capture microbes but assumes phenotypic and functional features of antigen-presenting cells, including the capacity to cross-present exogenous antigens to CD8 T cells. These findings indicate that steady-state mammalian NM harbors a unique innate cellular immune environment that is unlike any other barrier tissue.

## INTRODUCTION

The average adult human inhales ∼11,000 liters of air per day, mostly via the nasal passage^1^. As air travels through the nasal cavity (NC), it must be moistened and warmed (or cooled) before reaching the lower airways. Additionally, in most environments, air contains a myriad of organic and/or inorganic materials that must be removed to prevent potentially harmful effects on the lung^2,3^. Both the conditioning and filtering of inhaled air, as well as chemosensation of odors and flavorants, depend on the nasal mucosa (NM), a highly innervated epithelialized membrane that lines the NC and covers the underlying osseous and cartilaginous structures that shape the nasal passage. As air moves from the tip of the nose toward the nasopharynx (**Fig. 1A**), it first passes across stratified squamous epithelium, followed by progressively deeper anatomic regions that are lined by transitional (anterior maxiloturbinate), ciliated columnar (maxiloturbinate and nasoturbinate), and olfactory epithelial cells (posterior nasoturbinate and ethmoturbinate).

**Figure 1.**
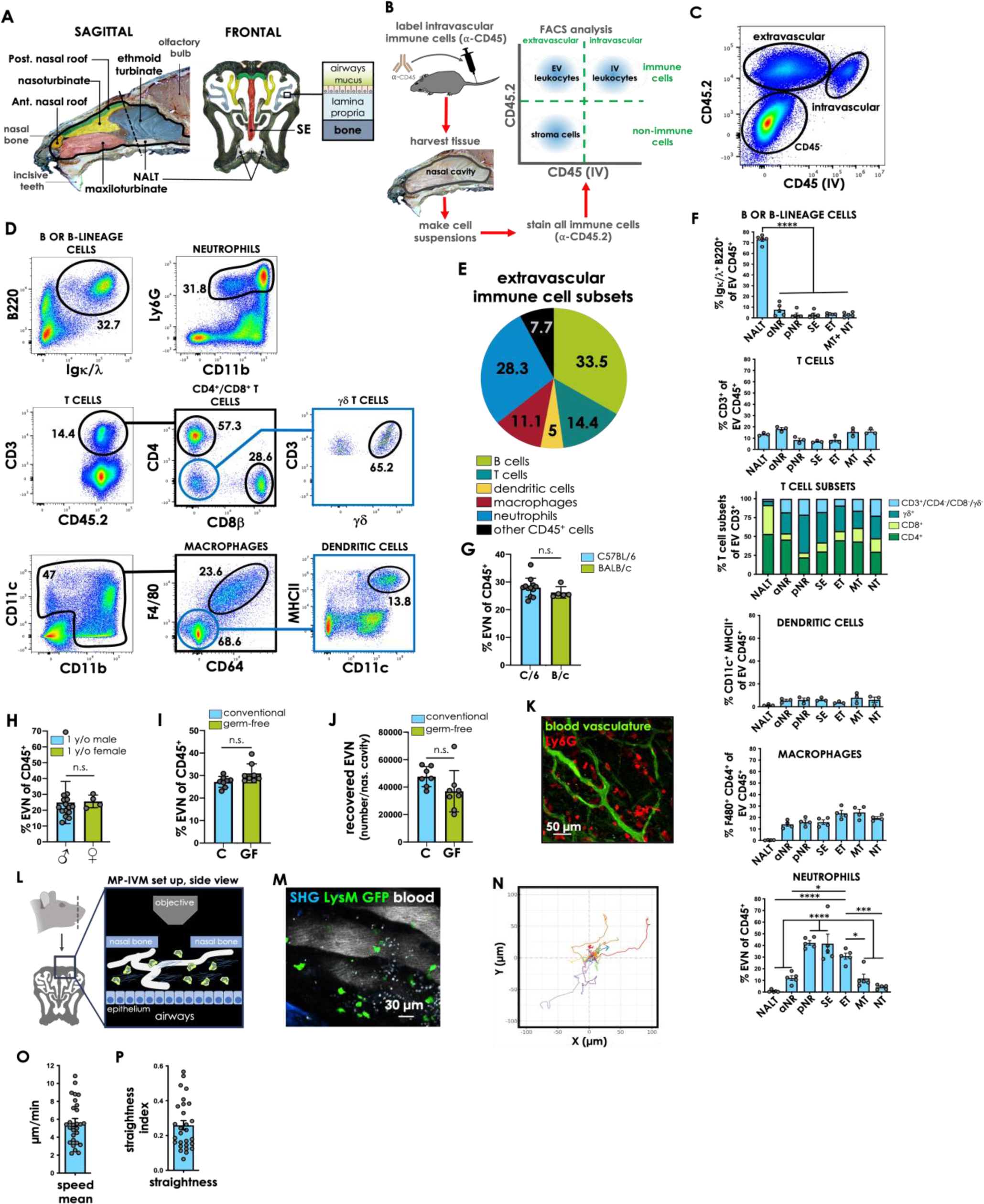
Extravascular neutrophils (EVN) in steady state NM. **(A)** Schematic of anatomic regions of the nasal cavity. Left: midline sagittal view. The nasal passage is demarcated by a black line with anatomic structures identified by color: anterior nasal roof (aNR, orange), posterior nasal roof (pNR, green), nasoturbinate (NT, yellow), ethmoid turbinate (ET, blue), maxiloturbinate (MT, pink), and nasopharynx-associated lymphatic tissue (NALT, gray). Middle: frontal view (section location corresponds to the dotted line in the sagittal view). Septum (SE, red) is shown only in the frontal view. Right: relative positions of bone, mucosa (consisting of lamina propria and epithelium), mucus layer, and airways. The mucosa in the anterior and posterior roof regions are accessible for IVM imaging. **(B)** Schematic of intravascular labeling strategy. **(C)** Representative dot plot resulting from the gating strategy shown in B. This strategy was used to identify extravascular leukocytes in single-cell suspensions of NM using FACS analysis. Gating on live singlets, based on live/dead staining. **(D)** Gating strategy to identify extravascular immune cell populations. **(E)** Frequency of different immune cell populations among extravascular CD45^+^ cells. Gating on the extravascular population shown in panel C. **(F)** Frequency of different immune cell populations among extravascular CD45^+^ cells by anatomical region. T cell subsets are shown as percentages of all T cells (CD3^+^). One-way ANOVA with Tukey’s multiple comparisons test. Differences between groups were considered significant when *P*<0.05. \**P*<0.05; \*\*\**P*<0.001; \*\*\*\**P*<0.0001. Data from one representative of two experiments. **(G)** Frequency of EVN among extravascular CD45^+^ cells in C57BL/6 (blue, C/6) and BALB/C (green, B/c) mice. Data are pooled from two independent experiments. **(H)** Frequency of EVN in NM of aged (≥1 year old) male (blue) and female (green) mice. Data are pooled from three independent experiments. **(I)** Frequency and **(J)** number of recovered EVN in the NC of conventional (blue, C) and germ-free (green, GF) mice. For **I** and **J**, data were pooled from two independent experiments. For **F-J**, circles represent a single mouse, bars represent the mean of the group, and error bars represent SEM. For **G-J**, the Mann-Whitney test was used and n.s denotes no significance. **(K)** Representative photomicrograph depicting EVN (red) in SE mucosa of a Ly6G-tdTomato mouse. The animal was injected IV with AF488-labeled wheat germ agglutinin (WGA) three minutes before euthanasia to label blood vasculature (green). **(L)** Schematic of 2P-IVM model depicting the anatomical site for imaging (left), and a side view of the tissue (right), with neutrophils (green), blood vasculature (white tubular structures), and collagen fibers (blue) in the lamina propria, flanked by the epithelium and the nasal bone. **(M)** Still image taken from Supplementary Video 1, showing LysM GFP neutrophils (green), blood vasculature (white), and bone (blue, lower left corner) demarcating a segment of the drilled burr hole’s edge. **(N)** Spider-plot of cell tracks from supplementary video 1. Each color represents a cell track from an individual cell, with the starting point of each track shifted to start from a common origin. **(O)** Neutrophil speed mean calculated from supplementary video 1. **(P)** straightness index, calculated from supplementary video 1. For O and P, the bar represents the mean value and the circles represent a tracked cell.

Most of the NM epithelium is covered by adhesive mucus that captures inhaled materials ^2,4–6^. Particles and debris that become trapped in this mucus are expelled from the NC by the action of beating cilia on the respiratory epithelium. By contrast, many air-borne pathogens have evolved mechanisms to avoid mucociliary clearance enabling them to instigate a local infection. Indeed, the NM is susceptible to infection by a multitude of viruses^7–10^, bacteria^11–13^, fungi^14^, and parasites^15^. Many of these pathogens pose a risk of disseminating into the lung or other tissues, such as the brain, middle ear, or blood, causing potentially life-threatening disease. In addition, the NC hosts a commensal microbiome and is also frequently colonized by pathobionts, opportunistic microorganisms that may cause infections under certain circumstances^13,16^. Given the multitude of microbial threats, it stands to reason that the NM must be well-guarded and capable of mounting rapid and potent immune responses upon invasion by diverse microorganisms.

In general, anti-microbial immune responses in barrier tissues are thought to be executed by the concerted action of immune cells that either reside within the tissue *a priori* or are recruited from the blood upon infection. However, we have a limited understanding of the immune landscape within the NM that is encountered by invading pathogens. Thus, we conducted a comprehensive survey of CD45^+^ immune cells that populate murine NM at steady state. Our analysis uncovered an unexpectedly large population of extravascular neutrophils (EVN) that expressed the diagnostic markers Ly6G and CD11b. Neutrophils are polymorphonuclear phagocytes that are continuously generated in BM and released into the blood circulation to monitor the body for acute infections or tissue injury^17^. Traditionally, neutrophils are thought to be short-lived, terminally differentiated cells that access peripheral tissues only when homeostasis is disrupted ^17,18^. However, recent studies suggest that distinct neutrophil subsets may exist that display diverse phenotypes (possibly reflecting different maturation stages) and may exert specialized functions^19,20^.

Here, we show that the steady-state NM of mice is constitutively populated by abundant EVN, even in the complete absence of microbial stimuli. Furthermore, our findings indicate that these nasal EVN are comprised of three phenotypically distinct subsets, each with a distinct origin. One population (nN1) expressed Ly6G and CD11b at intermediate levels indicating a less mature stage. Indeed, these cells localized primarily to BM cavities in close vicinity to the mucosal layer. A second, more mature EVN population (nN2) was CD11b^hi^Ly6G^hi^ and preferentially accessed the NM via the blood. This population, uniquely among EVN subsets, captured microbial pathogens introduced into the NC. After accessing the NM, nN2 EVN differentiated over ∼5 days to give rise to a third population (nN3) that expressed CD11c and SiglecF. nN3 EVN did not capture infectious microbes but displayed features of antigen-presenting cells (APCs), including the ability to cross-present exogenous antigens (Ags) to CD8 T cells. Our findings indicate that the mammalian NM harbors a vibrant immune landscape that is markedly distinct from other epithelialized barrier tissues and may have evolved to cope with the unique physical, chemical, and microbial challenges encountered in the uppermost respiratory tract.

## RESULTS

### The nasal cavity harbors a large population of extravascular neutrophils

To map the cellular immune landscape of the NM, we initially generated single-cell suspensions of undisturbed NM from young adult C57BL6/J mice that were harvested ∼3 min after IV injection of a fluorescent MAb against CD45, a pan-leukocyte marker (**Fig. 1B**). In this protocol, the IV injected MAb labels all CD45^+^ cells in contact with flowing blood but does not stain extravascular immune cells, which were identified *ex vivo* by staining cell suspensions with a second MAb against CD45.2 (**Fig. 1C**)^21^.

Concomitant staining for a panel of leukocyte lineage markers was used to analyze the cellular composition of the extravascular immune compartment using flow cytometry (**Fig. 1D, E**). Of note, this initial analysis was focused on regions of the NC that are continuously exposed to inhaled airflow.

Therefore, we did not include nearby tissues that possess distinct physiological functions, such as the vomeronasal organ^2,22^, or the lining of the maxillary sinus^23^. Notwithstanding, although our single-cell preparations for flow cytometry were primarily derived from mucosal soft tissue, they also contained cells residing within airway mucus as well as bone from the ethmoturbinate (ET) region. Bony structures from other nasal regions, as well as cartilage from the septum (SE), were excluded.

FACS analysis of extravascular leukocytes (i.e. cells that were not stained by IV anti-CD45, but bound *ex vivo* anti-CD45.2) identified diverse myeloid and lymphoid cell populations (**Fig. 1E and F**), including an unexpectedly large fraction of cells that expressed two typical neutrophil markers, Ly6G and CD11b. The NC is traditionally subdivided into five anatomically defined regions: nasoturbinate (NT), maxiloturbinate (MT), SE, ET, and nasopharynx-associated lymphatic tissue (NALT) ^2^ (**Fig. 1A**). In addition, we identified two regions forming the roof of the NC, immediately below the nasal bone, that each possessed characteristic morphologic features that rendered them distinct from each other and from the adjacent nasoturbinate (**Supplementary Fig. 1A and B**). The anterior nasal roof (aNR) region is a thin transparent membrane close to the tip of the nose that was characterized by a dense network of anastomosing capillaries, broad, irregularly shaped lymphatics, sparse peripheral innervation, and patchy ciliated (βIII-tubulin^+^) epithelial cells. By contrast, the posterior nasal roof (pNR) region is relatively thick and macroscopically opaque with prominent innervation, a uniform network of relatively narrow lymph vessels, and sagittal bundles of parallel microvessels with blood flowing in a caudal-rostral direction. EVN were present in all regions of the NC but their frequency among extravascular CD45^+^ cells was significantly higher in pNR, SE, and ET than in the other NM regions (**Fig. 1F**).

Because EVN frequency in the NALT was significantly low, we did not include this region in this study. Moreover, EVN were similarly abundant in the NM of young adult C57BL/6 and BALB/c mice (**Fig. 1G**), as well as in NM of male and female aged mice (**Fig. 1H**). In all NM samples, EVN consistently accounted for ∼25-35% of all extravascular immune cells. Because the NC hosts a diverse community of microorganisms^16^, we asked whether the unexpected abundance of EVN in NM was a response to commensal microbes or a subclinical infection. However, EVN percentages and total numbers were comparable between NM of conventionally housed and germ-free mice (**Fig. 1I, J**), indicating that EVN are not elicited by microbial signals but constitutively present in steady state NC. Consistent with these findings, a disseminated population of EVN was readily detectable in whole-mount preparations of NM obtained from unmanipulated reporter mice^24^ (**Fig. 1K**)

Furthermore, because neutrophils are traditionally described as highly motile cells^25^, we developed a multi-photon intravital microscopy (MP-IVM) model to visualize EVN movement within the NM (**Fig. 1L**). In our model, a burr hole was drilled in the nasal bone to expose the NM (aNR or pNR). In addition to the implementation of an in-house-built stereotactic stage to place the tissue perpendicular to the microscope’s objective, a heated metal ring was placed around the imaging area for temperature control. Precise temperature monitoring of the tissue was possible through a thermal probe located immediately adjacent to the imaged area. In line with previous observations on conventional neutrophils, EVN displayed high motility (**Supplementary Video 1 and Fig. 1 M, and O**). In aggregate, our analysis suggests that EVN undergo random walk-like motility at kinetics reminiscent of those described for lymphocytes in peripheral lymph nodes^26^. Contrary to the directed movement that is typically observed in neutrophils during recruitment due to injury or infection^26^, the random walk of EVN indicates undirected immune surveillance of steady-state NM (**Fig. 1N-P**).

### Phenotypic diversity of nasal EVN

Phenotypically, blood-borne neutrophils represent a single population with a uniform expression of Ly6G and CD11b (**Fig. 2A**)^27,28^. In contrast, our analysis of nasal EVN revealed two distinct populations – a CD11b^int^Ly6G^int^ fraction reminiscent of blood neutrophils, and a CD11b^hi^Ly6G^hi^ fraction, which was not found in blood (**Fig. 2B** and **Supplemental Fig. 2A**). CD11b^hi^Ly6G^hi^ EVN were also present in other mucosal tissues, such as NALT and the uterus (**Fig. 2C**). In contrast, secondary lymphoid tissues including the spleen and cervical lymph nodes (LN) which drain the NC contained only CD11b^int^Ly6G^int^ neutrophils. A closer phenotypic analysis of CD11b^hi^Ly6G^hi^ EVN in NM revealed that this population can be further subdivided into two subsets based on the presence or absence of CD11c and sialic acid-binding immunoglobulin-type lectin F (Siglec F), markers that are typically not expressed on neutrophils (**Fig. 2B, D**). Rather, CD11c (αX integrin) is primarily considered a marker of dendritic cells (DCs)^29^ and some macrophage subsets^30^, while Siglec F is preferentially expressed by eosinophils^31^. Overall, our data indicate that nasal EVN display unusual phenotypic diversity in NM, comprising three distinct subsets: CD11b^int^ Ly6G^int^ CD11c^-^ SiglecF^-^, CD11b^hi^ Ly6G^hi^ CD11c^-^ SiglecF^-^, and CD11b^hi^ Ly6G^hi^ CD11c^+^ SiglecF^+^. We will henceforth refer to these subsets as nasal neutrophils 1 (nN1), nasal neutrophils 2 (nN2), and nasal neutrophils 3 (nN3), respectively.

**Figure 2.**
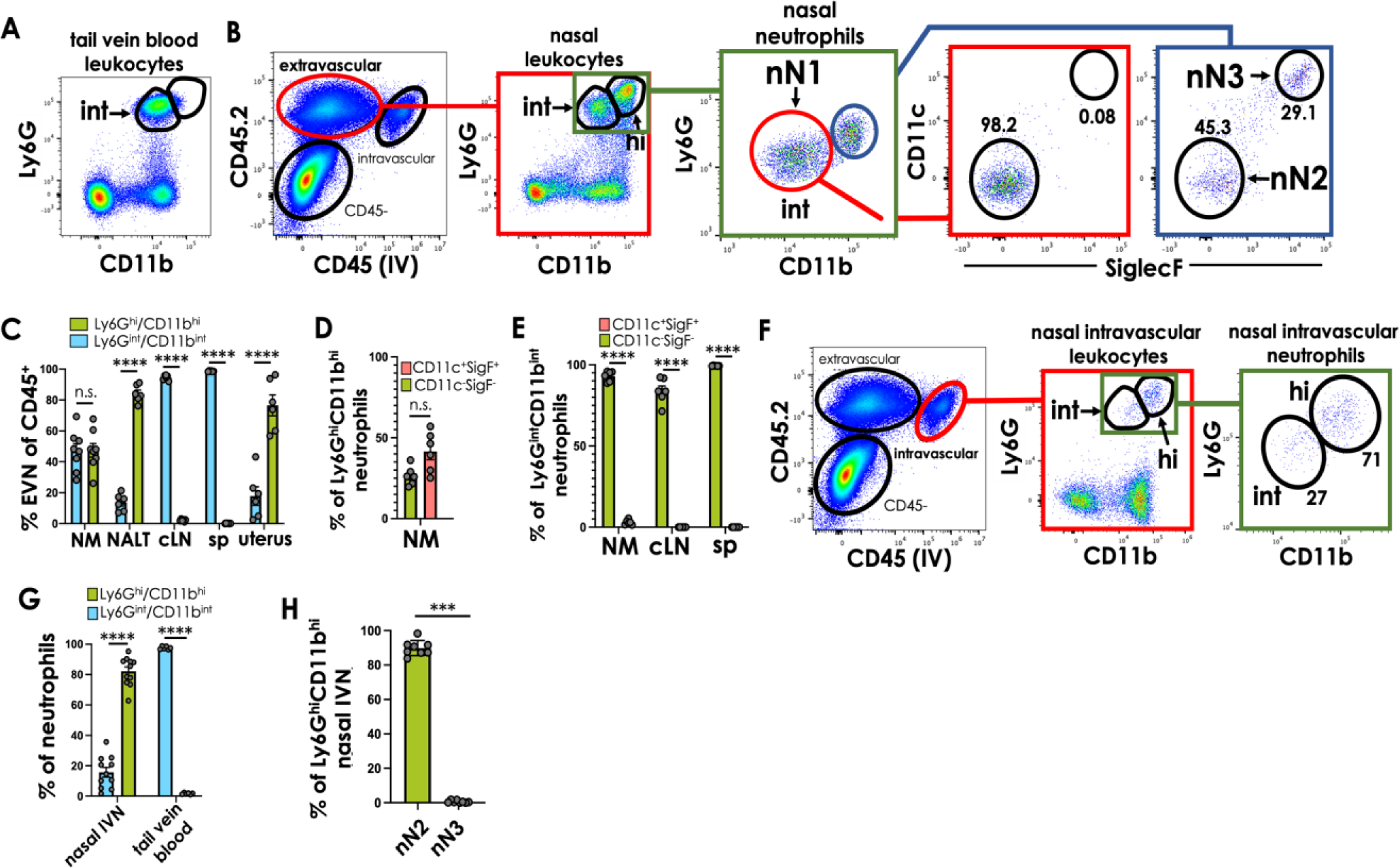
Nasal EVN constitute three phenotypically distinct subsets. (**A-B**) Representative FACS plots showing (**A**) blood neutrophils (gated on CD45^+^ cells) and (**B**) EVN in NM single-cell suspensions. Staining of classic neutrophil markers, Ly6G and CD11b, reveals two EVN populations expressing these markers at intermediate (nN1) or high levels. Ly6G^hi^CD11b^hi^ cells are further subdivided based on the absence (nN2) or presence of CD11c and SiglecF (nN3). (**C**) Frequency of CD11b^int^Ly6G^int^ and CD11b^hi^Ly6G^hi^ neutrophils in NM, nasal-associated lymphoid tissue (NALT), spleen (sp), and uterus. (**D**) Frequencies of nN2 and nN3 EVN in NM. (**E**) Frequencies of CD11c^-^SiglecF^-^ and CD11c^+^SiglecF^+^ neutrophils among CD11b^int^Ly6G^int^ neutrophils in NM and lymphoid tissues. (**F**) Representative dot plots showing CD11b and Ly6G expression of NM IVN. (**G**) Frequency of CD11b^int^Ly6G^int^ and CD11b^hi^Ly6G^hi^ among NM IVN and blood neutrophils. **(H)** Frequency of CD11c^-^SiglecF^-^ (nN2) and CD11c^+^SiglecF^+^ (nN3) neutrophils of CD11b^hi^Ly6G^hi^ NM IVN. For **C-E**, **G**, and **H**, data are pooled from two independent experiments. Statistic test in **C**, **E**, and **G** was one-way ANOVA, and in **D and H** Mann-Whitney test. Differences between groups were considered significant when p<0.05. ***P<0.001, ****P<0.0001; n.s., not significant. Circles in bar graphs represent a single mouse; bars represent group means, and error bars represent SEM.

We also examined the intravascular neutrophil (IVN) population in NM. In contrast to the CD11b^int^Ly6G^int^ phenotype of freely circulating blood neutrophils (**Supplementary Fig. 2A**), IVN in NM were predominantly CD11b^hi^Ly6G^hi^CD11c^−^SigLec F^−^ (**Fig. 2F, G & H, and Supplementary Fig. 2B**). These observations suggest a sequence of events whereby circulating CD11b^int^Ly6G^int^ neutrophils attach within the NM microvasculature and upregulate CD11b and Ly6G before or during the process of extravasation into the surrounding tissue. In this context, it should be noted that CD11b can be stored intracellularly to translocate to the cell surface within minutes after neutrophil activation by various stimuli^32^. As opposed to CD11b, levels of Ly6G did not increase on neutrophils under LPS stimulation, suggesting that the surface phenotype of IVN reflects intravascular differentiation rather than merely a response to activating stimuli (**Supplementary Fig. 2C**).

### Nasal EVN subsets have similar morphologies but distinct transcriptional profiles and origins

Although co-expression of Ly6G and CD11b is widely considered to be diagnostic for neutrophils^27,28^, the unusual phenotypic appearance of NM EVN, especially the N3 subset, prompted us to perform a morphological analysis to rigorously assess the identity of each of the three EVN populations. By flow cytometry, all three EVN subsets, as well as peripheral blood neutrophils, displayed similar forward and side light scatter profiles, rough indicators of cellular size and granularity, respectively (**Fig. 3A**).

**Figure 3.**
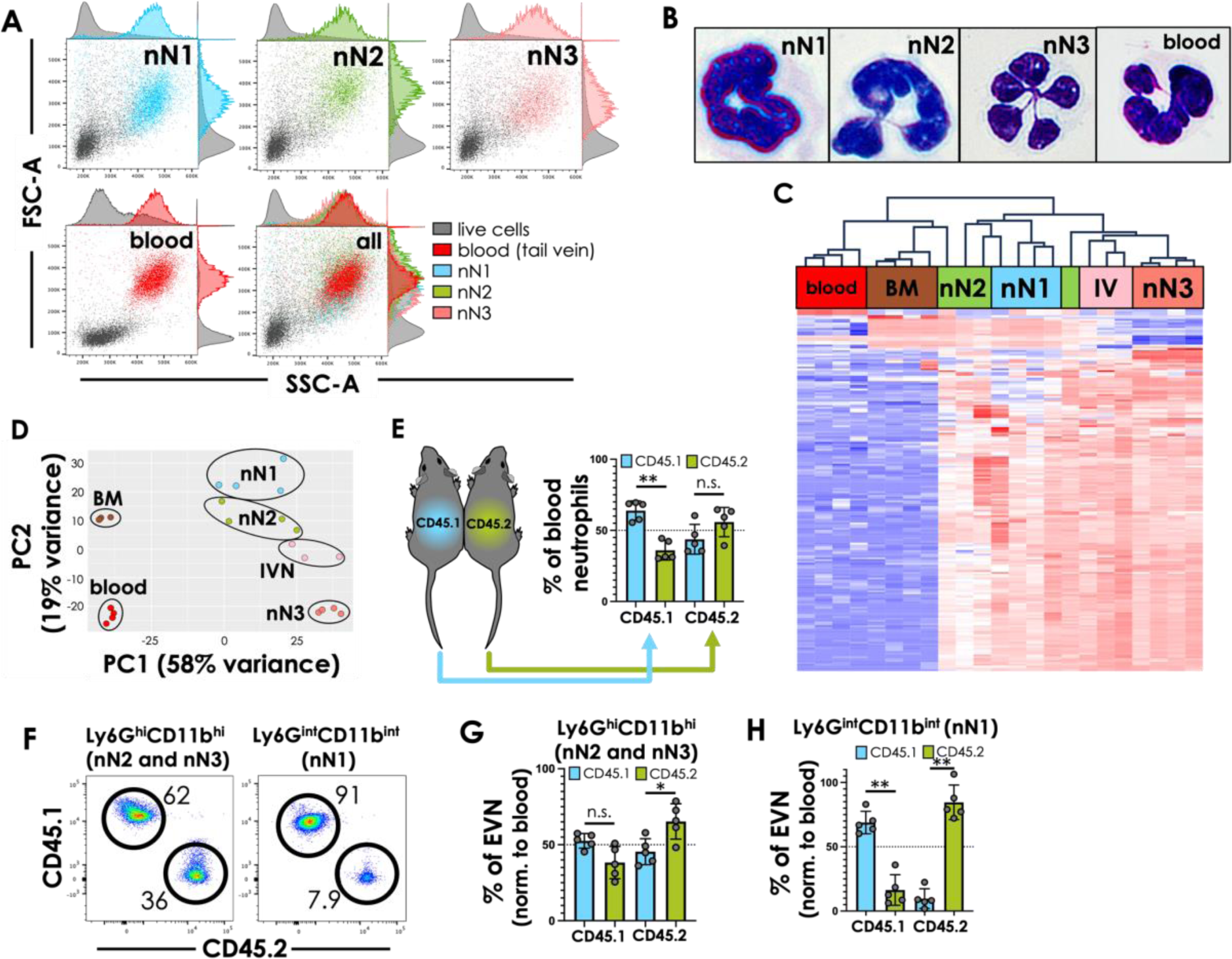
Differential origin of nasal EVN subsets. **(A)** Representative FACS dot plots showing forward and side scatter profiles of nN1, nN2, nN3, and blood neutrophils. Gray dots represent live immune and non-immune cells. **(B)** Representative micrographs of H&E stained cytospins of sorted nN1, nN2, nN3, and blood neutrophils. One representative of 10 similar images is shown for each subset. **(C)** Unsupervised hierarchical clustering of differential gene expression in neutrophils from blood, BM (femur), and NM IVN and EVN. Transcriptional profiles obtained by bulk RNAseq of FACS-sorted nN1, nN2, nN3, IVN, blood, and BM neutrophils. Normalized values of the 200 most variable genes shown, cut-off Z score of 4. Red: upregulated gene expression, blue: downregulated gene expression. **(D)** Principal component analysis (PCA) of mRNA transcription profiles obtained by RNAseq from FACS-sorted nN1, nN2, nN3, IVN, blood, and BM neutrophils. Each symbol represents a biological replicate. **(E)** Schematic of parabiosis (left) depicting a pair of congenic mice (CD45.1^+^ and CD45.2^+^) that were surgically conjoined at the flanks to share blood circulation. The bar graph (right) shows the frequency of host- and partner-derived neutrophils in blood of CD45.1^+^ and CD45.2^+^ parabionts (right) on week five of parabiosis. The dotted line depicts 50% chimerism. **(F)** Representative dot plots showing the frequency of CD45.1^+^ and CD45.2^+^ neutrophils among nN1, nN2, and nN3 EVN in a CD45.1^+^ parabiont. **(G-H)** Frequencies of host- and partner-derived EVN subsets in parabionts were assessed for (**G**) CD11b^hi^Ly6G^hi^ (nN2 and nN3) and (**H**) CD11b^int^Ly6G^int^ (nN1) EVN. Data in **E, G**, and **H** were pooled from two independent experiments and analyzed by Mann-Whitney test. Differences between groups were considered significant when p<0.05. *P<0.05; **P<0.01, n.s., not significant. Circles in bar graphs represent individual mice; bars and error bars represent group means and SEM, respectively.

Moreover, H&E staining of sorted EVN subsets revealed a characteristic polymorphonuclear appearance, whereby many nN1 cells showed a more band-like nuclear structure typical of young neutrophils, while the majority of N3 cells displayed hyper-segmented nuclei that are usually encountered in aged neutrophils (**Fig. 3B**)^27,33^. Notwithstanding, these findings support the conclusion that all three EVN populations are, in fact, phenotypically distinct subsets of neutrophils.

Next, to further explore the extent to which the three EVN populations in NM were similar or different from one another, we conducted a bulk RNA sequencing (RNAseq) analysis of FACS-sorted nasal EVN, as well as nasal IVN, and neutrophils from peripheral blood and femur bone marrow (BM).

Unsupervised hierarchical clustering of differentially expressed (DE) genes of the four nasal subsets (nN1, nN2, nN3, and IVN) and that of non-mucosal (BM and blood) neutrophils revealed that these populations are transcriptionally distinct (**Fig. 3C**). In addition, a principal component analysis (PCA) also indicated that each nasal EVN population is transcriptionally distinct from all other subsets, with the N3 neutrophil population being the furthest removed within the PCA space (**Fig. 3D and Supplementary table 1**).

### nN1 neutrophils originate within BM cavities adjacent to NM

Having confirmed the identity of NM EVN as neutrophils, we set out to investigate each subset’s origin. Because neutrophils usually access peripheral tissues via the blood circulation^17^, we asked whether EVN in NM also derive from circulating blood neutrophils. Thus, we conducted parabiosis experiments whereby pairs of congenic mice (CD45.1^+^ and CD45.2^+^) were surgically conjoined to establish a shared blood circulation for five weeks (**Fig. 3E**). Following this equilibration period, the chimerism of Ly6G^hi^CD11b^hi^ EVN (nN2 and nN3) was slightly lower than in blood, but nonetheless substantial, indicating that the nN2 and N3 subsets arise predominantly from migratory circulating neutrophils (**Fig. 3F and G**). By contrast, Ly6G^int^CD11b^int^ (nN1) EVN exhibited minimal chimerism in parabiotic mice (**Fig. 3F and H**).

In aggregate, the observation that the nN1 EVN subset did not access the NM from the circulation and displayed a less mature nuclear morphology and surface phenotype implies that nN1 EVN originate either within the NM or in its close vicinity. Indeed, in most regions of the NC, the NM tightly covers bony structures of the skull some of which contained bone marrow (BM) cavities (**Supplementary Fig. 3A**). Accordingly, in LysM GFP mice in which neutrophils selectively express a GFP reporter^34^, BM cavities within submucosal bone adjacent to NM were filled with GFP^bright^ cells, especially in the nasal bone and ET region (**Supplementary Fig. 3B-D**).

In our standard protocol to harvest NM from mice, the soft mucosal and submucosal tissue can be readily separated from underlying bones in most regions of the NC, except in the ET where the mucosa is tightly adherent to delicate boney structures and difficult to isolate without painstaking microsurgical dissection. Thus, in the experiments above, leukocytes isolated from non-ET regions were from pure mucosal samples, whereas ET cells represented a mixture of cells from either mucosa or bone/BM that were harvested together. To ask whether nN1 EVN reflect BM-resident cells, we analyzed cell suspensions that were individually isolated from aNR, pNR, NT, MT, NALT, SE, or ET regions. nN1 neutrophils represented the majority of EVN in ET but only amounted to ∼5-10% of EVN in all other regions (**Fig. 4A and B**). Moreover, after careful microsurgical separation of ethmoid bone structures and soft mucosal tissue (**Supplementary Fig. 3A**), nN1 neutrophils represented the vast majority of neutrophils in cell suspensions obtained from ET bone, whereas EVN in bone-free ET mucosa contained only ∼5% nN1 cells similar to other NM regions (**Fig. 4C and D**).

**Figure 4.**
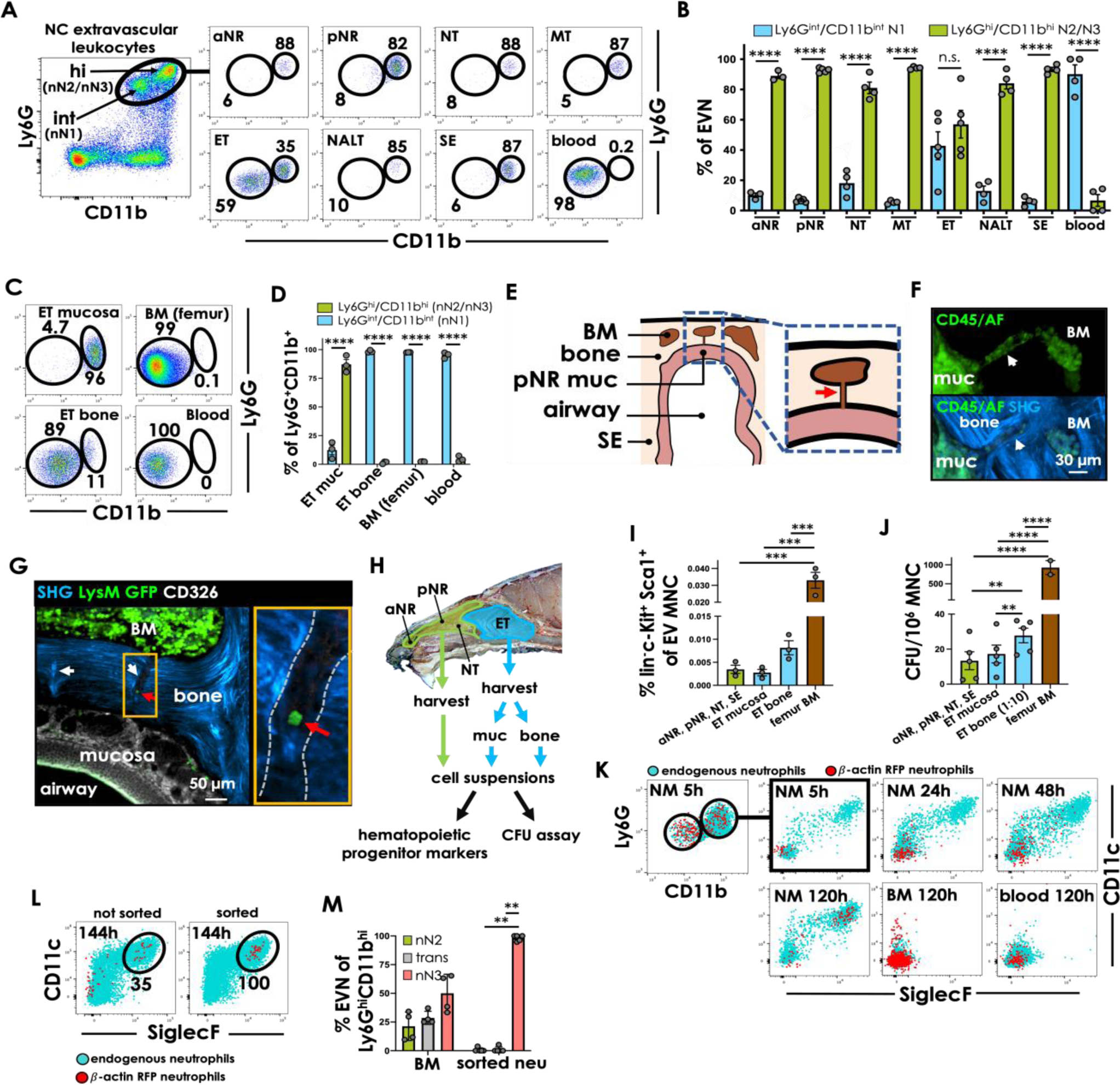
nN1 EVN reside in nasal BM, whereas nN2 EVN locate to NM and give rise to nN3 EVN. (**A**) Representative FACS plots depicting nN1 (CD11b^int^Ly6G^int^) and nN2/nN3 (CD11b^hi^Ly6G^hi^) EVN in different regions of the nose. Blood neutrophils are shown for comparison. **(B)** N1 (CD11b^int^Ly6G^int^) and nN2/nN3 (CD11b^hi^Ly6G^hi^) neutrophils in the different regions of the NC. nN1 neutrophils are present in the ET and largely absent in aNR, pNR, NT, MT, ET, NALT, and SE. Blood is shown for reference. Data pooled from two independent experiments. (**C**) FACS plots depicting nN1 (CD11b^int^Ly6G^int^) and nN2/nN3 (CD11b^hi^Ly6G^hi^) EVN in ET mucosa, ET bone, femur BM, and blood. Same gating strategy as in **A**. (**D**) Frequency of nN1 (CD11b^int^Ly6G^int^) and nN2/nN3 (CD11b^hi^Ly6G^hi^) EVN in ET mucosa, ET bone, femur BM, and blood. Data from two pooled experiments. (**E**) Schematic illustration of anatomic features visualized in panels **F** and **G** showing a frontal view of the left pNR region. The nasal bone (beige) contains cavities (brown) filled with BM that connect to the nearby mucosa (pink) via channel-like conduits (arrow). (**F**) Representative multi-photon micrograph depicting CD45^+^ cells labeled green and a conduit (arrow) connecting BM and mucosa (muc). The upper panel shows green fluorescence alone, emitted from CD45^+^ cells stained with a fluorescently labeled antibody, in addition to low levels of autofluorescence (AF). The lower panel shows green fluorescence and second harmonics generation (SHG) from bone (blue). (**G**) Representative multi-photon micrograph depicting BM, pNR mucosa, and two connecting conduits (white arrows) from a LysM-GFP mouse. The image on the right shows a magnified region, delineated by an orange rectangle, depicting a neutrophil (green, red arrow) within a conduit. **(H)** Schematic protocol for assessment of hematopoietic progenitors in different NM regions. **(I)** Frequency of CD45^+^ lineage^-^ (i.e.: CD3^-^, CD11b^-^, CD11c^-^, CD19^-^, Ly6G^-^, NK-1.1^-^, NKp46^-^, and Ter-119^-^), c-Kit^+^, and Sca1^+^ cells within indicated nasal tissues. Femur BM was used as a control. Bars represent mean ± SEM of progenitor frequency among extravascular mononuclear cells (EV MNC). (**J**) CFU counts (CFU/10^6^ MNC) in methylcellulose assays using culture conditions that allow the proliferation of MG-HP. Bars represent the median CFU/10^6^ MNC per group. For **B, D, I,** and **J**, circles represent samples from individual mice, and error bars represent SEM. Differences between groups were tested by one-way ANOVA and considered significant when p<0.05. **P<0.01; ***P<0.001; ****P<0.0001. For I and J, data from one representative experiment, out of three experiments, are shown. **(K)** Representative dot plot depicting endogenous EVN (blue) and donor β-actin RFP neutrophils (red) in the NM, at different time points after adoptive transfer into WT mice. (**L-M**) Representative dot plot (**L**) and quantification (**M**) comparing the phenotype of adoptively transferred unsorted BM cells (left) *versus* FACS-sorted neutrophils (right) from β-actin RFP donors at 144h post transfer. CD11c^low^ SiglecF^low^ neutrophils were considered transitioning (trans, gray bar in **M**). Data were pooled from two independent experiments and compared by Mann-Whitney test. Differences between groups were considered significant when p<0.05. **P<0.01. Circles represent individual mice; bars represent the mean ± SEM.

To better understand the spatial relationship of EVN, BM cavities, and adjacent mucosa in the NC, we imaged ∼100 µm thick frontal frozen sections of the pNR region from LysM^GFP^ mice (**Supplementary Fig. 3C**). By using two-photon microscopy (2PM), the bone matrix of the nasal bone can be readily visualized in unstained sections owing to a bright second harmonic generation (SHG) signal emitted by collagen fibers ^35^. This imaging strategy revealed that BM and NM are locally connected through channel-like structures embedded within the otherwise uniformly dense bone matrix (**Fig. 4E-G**). These channels were reminiscent of the bone conduits that were recently reported to connect calvarian BM and the meninges^36,37^. Of note, many nasal bone conduits contained CD45^+^ immune cells (**Fig. 4F**) including neutrophils (**Fig. 4G**), suggesting that at least some leukocytes that arise within BM adjacent to the NM may be able to directly access the NM via local channels rather than via the blood stream.

The existence of bone conduits in the NC could potentially explain the consistent presence of low, but detectable numbers of nN1 EVN throughout the NM (**Fig. 4A and B**). Two non-exclusive scenarios appear plausible: nN1 cells could either arise within BM cavities and then use bone conduits to migrate into the surrounding NM and/or they may be generated within the NM itself by migratory BM-derived hematopoietic stem or progenitor cells (HSPC). In support of the latter idea, flow cytometric analysis of mucosal samples that were either pooled from aNR, pNR, NT, and SE or isolated from bone-free ET preparations contained small numbers of CD45^+^ leukocytes that displayed an HSPC phenotype (lineage^-^ c-Kit^+^ Sca-1^+^ ^38^ ^39,40^; **Fig. 4H, I**, and **Supplementary Fig. 3F**). To further explore whether the NM harbors true granulocyte progenitors, we cultured NM cell suspensions in semisolid methylcellulose-based medium with cytokines that specifically promote colony formation by macrophage/granulocyte hematopoietic progenitors. Indeed, colony-forming units (CFU) were readily detectable in NM samples, albeit at a lower frequency than in BM (**Fig. 4J**). Of note, colonies contained cells of two discernible sizes consistent with the formation of macrophages and granulocytes, with the latter being smaller than the former^41^ (**Supplementary Fig, 3G**). These findings indicate that hematopoietic progenitors that possess the potential to generate neutrophils and other myeloid leukocytes access the NM, presumably via local bone conduits.

### After homing to the NM, nN2 neutrophils differentiate into nN3 neutrophils

Although our parabiosis experiments indicate that most nN2 and nN3 EVN derive from circulating cells, we noted that the hallmark markers of the nN3 subset, CD11c and SiglecF, are not expressed on either blood-borne neutrophils which present an nN1-like phenotype (**Fig. 2G & Supplementary Fig. 2A**) or on nasal IVN that apparently upregulated Ly6G and CD11b prior to diapedesing to join the nN2 EVN pool (**Fig. 2F-H & Supplementary Fig. 2B**). Thus, we hypothesized that circulating neutrophils that adhere within NM microvessels differentiate first into nN2 neutrophils and, after diapedesing, into nN3 neutrophils. To test this, we adoptively transferred to WT recipients BM mononuclear cells that contained ∼60% neutrophils (CD11b^int^ Ly6G^int^ CD11c^-^ SiglecF^-^) and expressed red fluorescent protein (RFP) under the β-actin promoter (**Supplementary Fig. 3H**). 5h after transfer, extravascular RFP^+^ neutrophils displaying the nN2 phenotype (CD11b^hi^Ly6G^hi^CD11c^-^SiglecF^-^) were readily detectable in the NM. These newly homed cells were initially CD11c and SiglecF negative, but a modest increase in CD11c was observed in some homed cells starting at the 24-hour time point. A few scattered donor EVN began to display a full-fledged nN3 phenotype by 48h, and the majority of the transferred neutrophils had become CD11c^+^ SiglecF^+^ by 120h (**Fig. 4K**). Notably, this phenotypic change of transferred neutrophils occurred exclusively in the NM; donor neutrophils that persisted in recipient blood or BM remained unchanged (**Fig. 4K**).

To rigorously rule out that a non-neutrophil precursor might give rise to nN3 neutrophils, we FACS sorted CD11b^int^Ly6G^int^CD11c^-^SiglecF^-^ neutrophils from BM of β-actin RFP expressing donor mice and injected them IV into WT recipients. At six days after adoptive transfer, nearly 100% of the donor neutrophils within recipient NM had adopted the nN3 phenotype (**Fig. 4L and M**). Taken together, these results indicate that blood neutrophils, upon adhering within NM microvessels adopt an nN2 phenotype and, after emigrating into the NM, differentiate into nN3 neutrophils, a process that may last up to 6 days.

### nN2 neutrophils preferentially interact with pathogens in the NC

Having clarified the origins of each nasal EVN subset, we next sought to explore their functional properties. Because neutrophils are generally thought of as the first line of defense against invading bacteria^42^, we examined the response of nasal EVN to IN challenge with *Streptococcus pneumoniae*, a bacterial pathogen that frequently causes respiratory tract infections^43,44^. To facilitate bacterial detection and tracking, we used a genetically modified strain of *S. pneumoniae* that constitutively expressed green fluorescent protein (GFP). To induce a localized infection within the NC, 1×10^7^ CFU of fluorescent *S. pneumoniae* were instilled IN in 5 µL PBS per nostril. This inoculation protocol was based on pilot experiments indicating that an inoculum volume of up to 5µl remains confined to the NC and does not ‘spill over’ to the lower airways. Fifteen hours after challenge, NM was harvested and each EVN subset was analyzed by flow cytometry and fluorescence microscopy to assess their association with GFP^+^ bacteria.

There was minimal association between nN1 EVN and *S. pneumoniae*, which was not unexpected given the sheltered localization of most nN1 EVN within BM cavities. By contrast, although both nN2 and nN3 EVN reside in the NM, GFP^+^ bacteria were almost exclusively detected in association with nN2 EVN, whereas nearly all nN3 EVN remained GFP negative (**Fig. 5A and B**). *Ex vivo* microscopy of isolated nN2 EVN revealed that the bacteria localized intracellularly, indicating that nN2 EVN efficiently phagocytose invading microbes (**Fig. 5C**). To account for the striking paucity of *S. pneumoniae* in nN3 EVN, we considered the possibility that these cells may have lost the capacity to engulf bacteria, perhaps as a result of senescence. To test this idea, FACS purified EVN subsets were incubated for three hours with *S. pneumoniae in vitro*. In stark contrast to our *in vivo* observations, all three EVN subsets phagocytosed *S. pneumoniae* comparably (**Fig. 5D,E**, and **Supplementary Fig. 4A**), indicating that both nN1 and nN3 EVN are not inherently dysfunctional. To ask whether the lack of nN3 responsiveness *in vivo* reflects a specific inability of nN3 cells to respond to *S. pneumoniae*, a gram-positive extracellular bacterium, we tested neutrophil responses to two other phylogenetically and physiologically distinct pathogens that can colonize and/or infect the respiratory tract - *Chlamydia muridarum*, an obligate intracellular Gram-negative bacterium^45^, and *Candida albicans*, an opportunistic fungus. IN inoculation of genetically modified *C. muridarum* expressing a red fluorescent protein (dsRed) resulted in the formation of inclusions inside epithelial cells, confirming productive infection in the NM (**Supplementary Fig. 4B**). Neutrophils were found to engage in close interactions with free *Chlamydia* bacteria in the airways (**Supplementary video 2** and **Supplementary Fig. 4C and D**) and with epithelial cells containing *C. muridarum* inclusions (**Figure 5F** and **Supplementary video 3**).

**Figure 5.**
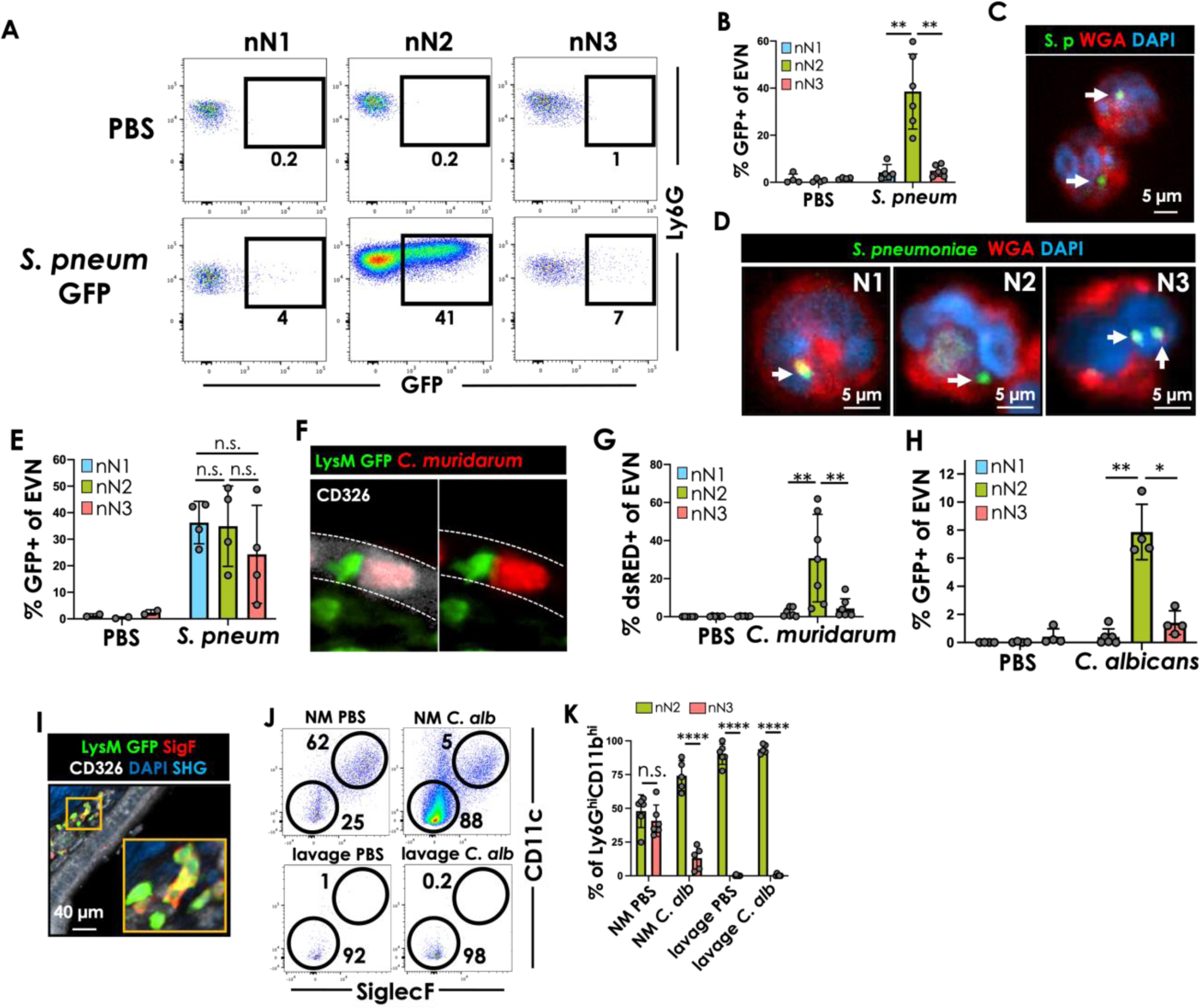
Only nN2 neutrophils access pathogens in the NC. (**A**) Representative dot plots of nN1, nN2, and nN3 neutrophils 15 h after intranasal instillation of PBS or *S. pneumoniae*-GFP. (**B**) Frequencies of GFP^+^ nN1, nN2, and nN3 neutrophils after PBS or *S. pneumoniae-*GFP instillation into the NC. Data are pooled from two independent experiments with five mice each. (**C**) Micrograph of sorted nN2 neutrophils from NM of a mouse that had been inoculated 15h earlier with *S. pneumoniae-* GFP (S.p., green). Cell nuclei and surface were stained with DAPI (blue) and wheat germ agglutinin (WGA, red), respectively. (**D**) Confocal micrographs of sorted nN1, nN2, and nN3 neutrophils 3h after *in vitro* incubation with *S. pneumoniae-*GFP (arrows; multiplicity of infection 0.1). For **C** and **D**, one representative of 10 similar micrographs is shown for each EVN subset. A single optical slice from a stack is shown, demonstrating the bacteria are inside the cell. (**E**) Flow cytometric quantification of the experiment described in **D** showing frequencies of GFP^+^ nN1, nN2, and nN3 neutrophils after *in vitro* exposure of sorted EVN subsets to PBS or *S. pneumoniae-*GFP. Pooled data from two independent experiments are shown. (**F**) Representative multi-photon micrograph of the NM of a LysM GFP mouse 48 h after intranasal instillation of *C. muridarum-*dsRed. CD326^+^ epithelium is shown in white, *C. muridarum* in red, and neutrophils in green. (**G**) Frequencies of dsRED^+^ nN1, nN2, and nN3 neutrophils 48h after PBS or *C. muridarum-*dsRED instillation into the NC. (**H**) Frequencies of GFP^+^ nN1, nN2, and nN3 neutrophils 15h after PBS or *C. albicans-*GFP instillation into the NC. (**I**) Representative multi-photon micrograph of neutrophils in the NM of a LysM GFP mouse after staining for SiglecF (red), epithelium (CD326, white), and cell nuclei (DAPI, blue). Bone is visualized by SHG in bright blue. nN2 and nN3 neutrophils appear in green and yellow, respectively. (**J**) Representative dot plots of nN2 (CD11c^-^ SigledF^-^), and nN3 (CD11c^+^ SigledF^+^) EVN in the NM or nasal lavage 15 h after instillation of PBS or *C. albicans* into the NC. (**K**) Frequencies of nN2 and nN3 EVN in the NM or nasal lavage 15 h after intranasal instillation of PBS or *C. albicans*. Data are pooled from two independent experiments with three mice each. For **B**, **E**, **G**, **H**, and **K**: Circles represent individual mice; bars represent group means and error bars represent SEM. Groups were compared by one-way ANOVA and differences between groups were considered significant when p<0.05. *P<0.05 **P<0.01 ****P<0.0001.

Analogous to our observations with *S. pneumoniae*, only nN2 EVN associated efficiently with *C. muridarum*, whereas interactions with nN1 or nN3 neutrophils were rarely detected (**Fig. 5G** and **Supplementary Fig. 4E**). Similarly, after IN inoculation of GFP-^46^ or mCherry-expressing^47^ *Candida albicans* only nN2 but not nN3 EVN were found to associate with this fungal pathogen (**Fig. 5H** and **Supplementary Fig. 4F and G**). These findings demonstrate that nN2 EVN are highly efficient at detecting and combating a variety of microbial pathogens in the NC, whereas the more differentiated nN3 subset does not participate in the performance of these classical neutrophil functions.

To further explore the striking difference in the ability of nN2 vs. nN3 EVN to engage with respiratory pathogens, NM sections from LysM GFP mice were stained with α-SiglecF MAb to visualize nN2 (GFP^+^) and nN3 (GFP^+^SiglecF^+^). Both populations resided in close proximity to each other within the NM (**Fig. 5I**), indicating that both EVN subsets should have similar access to microbes invading the subepithelial space. However, after introduction into the nasal airways, many pathogens may initially be trapped within mucus without traversing the epithelial barrier. To combat these microorganisms, EVN may need to access the infected airways. Indeed, analysis of nasal lavage fluid collected 15 hours after IN challenge with *C. albicans* revealed abundant nN2 neutrophils, whereas nN3 neutrophils were essentially absent (**Fig. 5J and K**), indicating that the apparent inability of the nN3 subset to engage with airway pathogens may be a consequence of their confinement to the subepithelial compartment.

### Transcriptome profiles of NM neutrophils indicate linear EVN subset differentiation

The multi-faceted distinctions in origin, phenotype, localization, function, and state of differentiation between the neutrophil subsets encountered within the NM prompted us to further compare these populations at the transcriptome level. As shown above, bulk RNAseq analysis of FACS-purified EVN, IVN, blood, and BM (femur) neutrophils revealed that these populations were different from each other at the transcriptomic level (**Fig. 3C and D**). To infer potential lineage relationships between EVN subsets, we performed single-cell RNA sequencing (scRNAseq) on neutrophil-enriched single-cell suspensions generated from all nasal regions, except NALT. These tissues were obtained from LysM GFP mice ∼3 min after IV injection of an oligonucleotide-barcoded MAb against CD45. The MAb binds to all intravascular immune cells, while extravascular immune cells remain unlabeled. Cell suspensions were subsequently labeled with oligonucleotide-barcoded MAb against Ly6G, CD11b, CD11c, and SiglecF, and neutrophils were isolated by FACS-sorting of GFP^bright^ cells (**Fig 6A**). RNA levels for each of the four markers used to distinguish EVN subsets were comparable in both bulk RNAseq (**Supplementary Fig. 5A**) and scRNAseq (**Fig. 6B**). After the exclusion of IVN (CD45^+^) cells from scRNAseq samples, we resolved seven cell clusters (**Fig. 6C**). Based on the measured protein expression, we ascribed clusters 3, 4, and 6 to the nN1 subset, clusters 0, 1, and 2 to the nN2 subset, and cluster 5 to the nN3 subset (**Fig. 6B, C and D**). To infer progenitor-progeny relationships between these clusters, we implemented a pseudotime analysis using Monocle 3, which learns the series of transcriptional changes that cells are undergoing and infers a trajectory^48^. Informed by our findings that nN1 EVN reflect mainly BM resident young neutrophils, we chose the trajectory origin within the clusters ascribed to nN1 EVN. The calculated pseudotime trajectory advanced from the nN1 EVN clusters into the clusters ascribed to nN2 EVN and ended in cluster 5, ascribed to the nN3 subset. Overall, this analysis is consistent with our experimental results outlined above and strongly implies a linear nN1->nN2->nN3 differentiation pathway.

**Figure 6.**
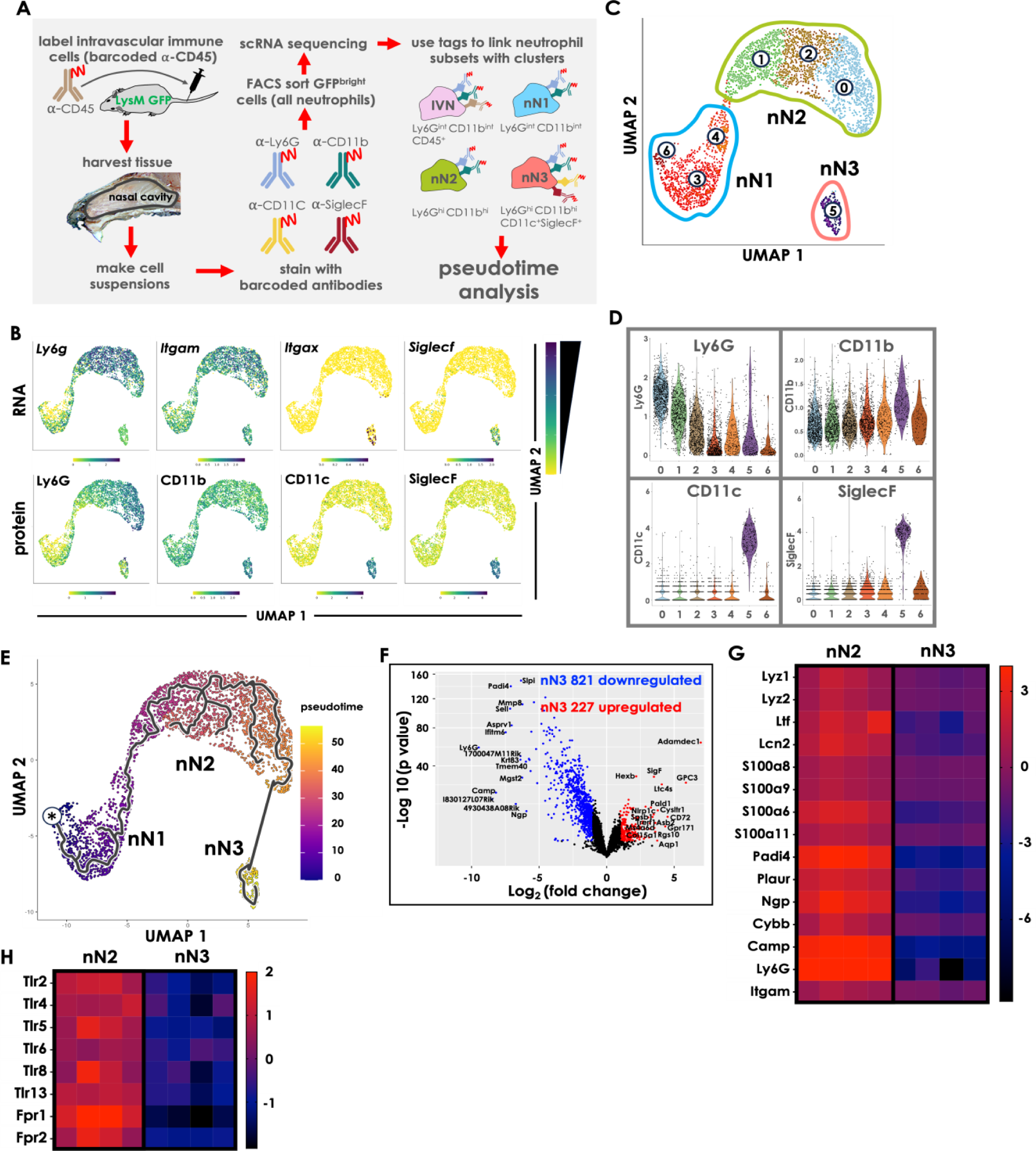
Transcriptome analysis of NM neutrophil subsets. (**A**) Schematic of scRNA sequencing strategy. (**B**) UMAP plot representation of EVN and their expression (RNA) of the genes *Ly6g*, *Itgam*, *Itgax*, and *Siglecf*, and their corresponding proteins Ly6G, CD11b, Cd11c, and SiglecF. (**C**) UMAP plot representation of EVN. Clusters that are ascribed to nN1, nN2, and nN3 EVN are demarcated with a blue, green, and pink line, respectively. (**D**) Violin plots indicating the expression distribution of the surface markers Ly6G, CD11b, Cd11c, and SiglecF in each cluster shown in C. (**E**) UMAP representation of EVN depicting pseudotime analysis (Monocle 3) with the green line representing cell developmental trajectory, and the asterisk representing the starting point of the trajectory. Lower values in the color scale represent early stages of the developmental trajectory. (**F**) Volcano plot displaying differentially expressed genes that are upregulated (red dots, 227 genes) or downregulated (blue dots, 821 genes) by nN3 EVN as compared to nN2 EVN. The top 15 most differentially up-or downregulated genes (by fold change) are identified. (**G**) Heatmap showing nN2 vs. nN3 EVN expression of 15 selected genes that are commonly ascribed to neutrophil functions. (**H**) Heatmap showing nN2 vs. nN3 EVN expression of 8 selected genes involved in neutrophil detection of microbial pathogens. For **G** and **G**, each column represents an independent biological replicate of sorted nN2 or nN3 EVN. Color corresponds to relative levels of expression within each row and the color scale sidebar numbers represent Z-score.

A more detailed comparison of nN2 vs. nN3 transcriptional profiles revealed that 227 genes were upregulated and 821 genes were downregulated (adjusted *p < 0.05* and absolute Log_2_ fold change > 1) in nN3 compared to nN2 EVN (**Fig. 6F**). Among the differentially downregulated genes were numerous genes associated with conventional neutrophil functions such as lysozyme, lactoferrin, lipocalin, and calprotectin (**Fig. 6G and Supplementary Fig. 5B**). Similarly, nN3 EVN had significantly downregulated genes that are essential for pathogen sensing, such as Toll-like receptors (TLRs)^49^ and the formyl peptide receptors, FPR1 and FPR2 ^50–52^ (**Fig. 6H and Supplementary Fig. 5B**). Accordingly, a Gene Ontology (GO) analysis of nN3 downregulated genes identified a set of genes involved in neutrophil degranulation and response to bacterial components (**Supplementary Table 2**).

### nN3 neutrophils exhibit features and functions of antigen-presenting cells

The observed loss of expression of classical neutrophil-associated genes in nN3 EVN implied a profound shift in identity and function of this subset. Indeed, a GO analysis of nN3 upregulated genes revealed a marked enrichment of genes involved in antigen processing and presentation (**Supplementary Table 3**). Furthermore, a comparative gene set scoring analysis, comparing differentially expressed genes in nN3 EVN to murine immune cell populations within the Immunological Genome Project (ImmGen^53^) database, found nN3-upregulated genes to be preferentially expressed in dendritic cells (DCs), macrophages, and monocytes, whereas nN3 downregulated genes were expressed by conventional neutrophils (**Fig. 7A)**.

**Figure 7.**
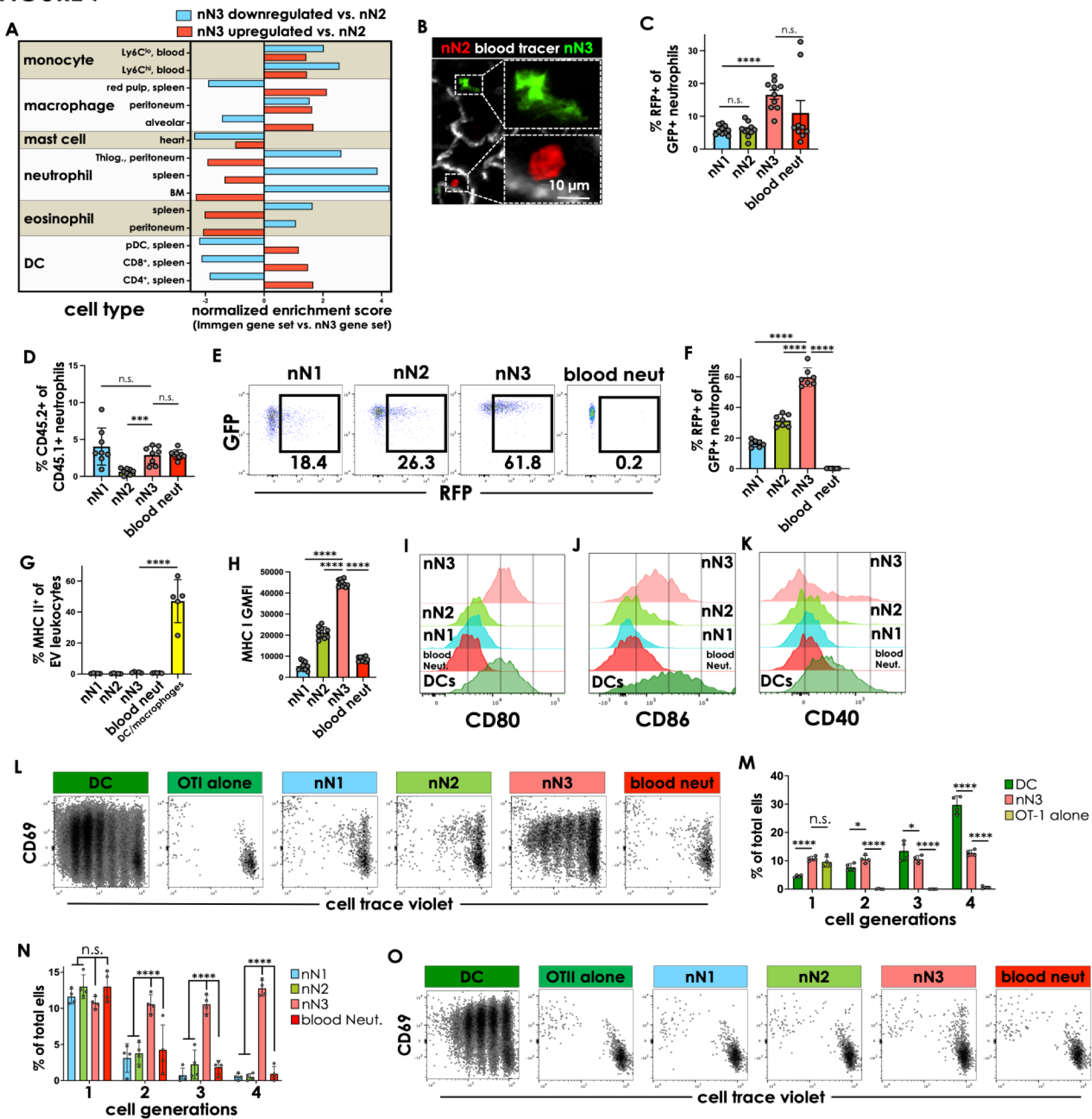
nN3 EVN are reprogrammed within NM to assume phenotypic and functional features of APC. **(A)** Enrichment scores of genes that were upregulated (red) or downregulated (blue) in nN3 EVN based on comparison to gene expression profiles from the Immunological Genome Project (ImmGen) in indicated leukocyte subsets and source tissues. **(B)** Representative multi-photon micrograph showing nN2 (red) and nN3 EVN (green) in an NM explant. See also Supplementary Figure 6A. (**C**) Frequency of GFP^+^ cells among β-actin-RFP nN1, nN2, and nN3 EVN in NM, and blood neutrophils. Samples were harvested from mixed BM chimeric WT hosts that had been lethally irradiated and transplanted with a 1:1 mixture of BM from β-actin-RFP and β-actin-GFP donors. (**D**) Frequency of phenotypically CD45.2^+^ cells among genetically CD45.1^+^ blood neutrophils as well as nN1, nN2, and nN3 EVN. Samples were harvested from lethally irradiated CD45.2^+^ mice that were transplanted with 1:1 BM from CD45.1^+^ and CD45.2^+^ donors. (**E,F**) GFP and RFP expression was assessed in blood neutrophils and in nN1, nN2, and nN3 EVN of lethally irradiated β-actin-RFP mice that had been transplanted with BM from β-actin-GFP donors. Panels show (**E**) representative FACS plots and (**F**) mean +/-SEM in n=7 animals. (**G**) Frequency of MHC-II^+^ cells in each neutrophil subset and in CD11c^+^ CD11b^+^ Ly6G^-^ cells. (**H**) Geometric mean fluorescence intensity (GMFI) of MHC-I staining on nN1, nN2, nN3 EVN, and blood neutrophil expression. (**I-K**) Representative histogram showing nN1, nN2, nN3, blood neutrophil, and DC expression of DC80 (**I**), CD86 (**J**), and CD40 (**K**). (**L**) Representative dot plots showing CD69 induction and proliferation of cell trace violet-labeled OT-I CD8 T cells after 72h incubation with sorted OVA-pulsed blood neutrophils, nN1, nN2, nN3 EVN, or Flt3L-induced BM DCs. (**M**) Quantification of OT-I cell proliferation (assessed by cell trace violet dilution) after co-culture with OVA-pulsed DC and nN3 EVN. OT-1 cells alone were used as negative control. (**N**) Quantification of OT-I cell proliferation (assessed by cell trace violet dilution) after co-culture with OVA-pulsed nN1, nN2, nN3, or blood neutrophils. For M and N, data are shown as the percentage of total cells of a cell subset, grouped by cell generation, for a total of four cell generations. (**O**) Representative dot plots showing CD69 induction and proliferation of cell trace violet-labeled OT-II CD4 T cells. Assay conditions were analogous to those in panel **L**. In all bar graphs, circles represent individual mice, bars represent group means and error bars represent SEM. Statistical differences between groups were assessed using one-way ANOVA (**C, D, F, H, M, and N**) or Mann-Whitney test (**G**) and considered significant when p<0.05. *P<0.05; **P<0.01, ***P<0.001; ****P<0.0001. Data from two pooled experiments are shown.

To further explore this observation at a morphological level, we took advantage of the fact that adoptively transferred neutrophils home to NM and uniformly differentiate into nN3 EVN within 5-6 days (**Fig. 4J and K**). Thus, naïve WT recipient mice received a first injection of β-actin-GFP neutrophils on day 0 and a second injection of β-actin-RFP neutrophils on day 5. This allowed for the unambiguous identification of nN2 and nN3 subsets within the NM on day 6 based on their distinct expression of RFP and GFP, respectively (**Supplementary Fig. 6A**). *In situ* imaging of fresh tissue explants revealed that nN2 and nN3 EVN were morphologically distinct, with nN2 neutrophils displaying a rounded, ameboid shape, as expected of conventional neutrophils, while nN3 neutrophils extended dendritic processes, resembling the appearance of DC^54^ (**Fig. 7B**).

In light of these conspicuous morphologic characteristics of nN3 cells and their apparent adoption of a gene expression profile typical of myeloid APC, we asked whether nN3 neutrophils possess the ability to acquire, process, and present exogenous antigens (Ag). Professional APC, such as DC and macrophages, can acquire antigenic material from other cells using diverse contact-dependent mechanisms^55–58^. Since nN3 EVN are in intimate contact with other immune cells in the NM (**Supplementary Fig. 6B**), we investigated if they could obtain material from surrounding hematopoietic cells. To this end, we generated mixed BM chimeric mice by transplanting a 1:5 mixture of BM cells from β-actin-GFP and β-actin-RFP donors into irradiated WT recipients, resulting in a corresponding ∼1:5 chimerism of GFP^+^:RFP^+^ neutrophils in peripheral blood (**Supplementary Fig. 6C**). In this setting, neutrophils are genetically restricted to express only one fluorescent protein, so any double positive cells would be indicative of having acquired material from one or more other hematopoietic cells.

FACS analysis of β-actin-GFP neutrophils in mixed BM chimeric mice revealed minimal RFP acquisition by blood neutrophils or nN1 and nN2 EVN in NM. By comparison, the frequency of double-positive cells was significantly higher, albeit still modest, in the nN3 subset (**Fig. 7C** and **Supplementary Fig. 6D**). Next, we asked whether nN3 EVN could also obtain and present surface proteins from neighboring leukocytes, a phenomenon sometimes referred to as ‘cross-dressing’^59^. To this end, a second set of BM chimeric mice was generated with hematopoietic cells expressing either the CD45.1 or CD45.2 surface markers (**Supplementary Fig. 6E**). In these animals, the frequency of double-positive neutrophils was negligible (**Fig. 7D** and **Supplementary Fig. 6F**). Lastly, we generated BM chimeric animals by transplanting β-actin-GFP BM into β-actin-RFP recipients to ask whether neutrophils acquire material from non-hematopoietic cells. In this setting, double-positive neutrophils were more frequent, particularly among nN3 EVN where the majority was robustly positive for RFP (**Fig. 7E and F**).

Having determined that nN3 neutrophils efficiently acquire potentially antigenic material from their environment, particularly from non-immune cells, we asked whether nN3 cells can present exogenous Ag to T cells. A number of studies have reported that certain neutrophil subsets can develop APC functions in response to a variety of stimuli^60^, however, neutrophils have not been shown to spontaneously acquire APC properties at steady-state. Moreover, the previously known modalities to induce APC functions in neutrophils usually result in robust expression of MHC-II, whereas NM EVN, including the nN3 subset, did not express detectable MHC-II on their surface or intracellularly (**Fig. 7G** and **Supplementary Fig. 6G, H and I**). By contrast, nN3 EVN expressed much higher levels of MHC-I than any other neutrophil subset in NM or blood (**Fig. 7H and Supplementary Fig. 6J-L**). nN3 EVN, but not nN1 or nN2 EVN, also expressed the costimulatory receptor, CD80, at levels comparable to NM-resident DCs (**Fig. 7I**). In addition, nN3 also expressed CD86 and CD40 at higher levels than the other subsets, albeit at lower levels than DC (**Fig. 7J and K**). This phenotype of nN3 EVN suggested that they might present Ag only in MHC-I to elicit a CD8 T cell response, a process known as cross-presentation that is traditionally considered a hallmark of type 1 conventional dendritic cells (cDC1)^61^.

To formally test if nN3 neutrophils can cross-present exogenous Ag to activate T cells, EVN subsets were isolated from NM and incubated with chicken ovalbumin (OVA) prior to co-culture with naïve OVA-specific CD8 or CD4 T-cells (OT-I and OT-II, respectively). nN1, nN2, and blood neutrophils failed to activate either T cell subset, however, OVA-pulsed nN3 EVN triggered robust CD8 T cell proliferation (**Fig. 7L, M and N**) with concomitant changes in the expression of T cell activation markers (**Supplementary Fig. 6M**). In contrast, and in agreement with their lack of MHC-II expression, nN3 EVN did not activate CD4 T cells (**Fig. 7O**). Of note, both DC and nN3, but not other EVN subsets also expressed robust levels of coinhibitory molecules including PDL-1 and TIM-3, whereas PDL-2 was only expressed on DC (**Supplementary Fig. 6 N-P**). Additionally, steady-state nN3 expressed negligible levels of instructive T-cell cytokines such as IL-12, IL-10, IL-4, IL-18, or INFγ (**Supplementary Fig. 6Q**).

## DISCUSSION

A crucial function of the NM is to retain inhaled particles to prevent pathogens from reaching deeper tissues (e.g. lungs and the brain). Consequently, the NM cellular immune responses are the gatekeepers that not only control local infection but also prevent its spread to the lungs and the brain. Nevertheless, the immune cell landscape of the NM and its role during homeostasis and infection remains understudied. Here, we investigated the cellular immune landscape of the NM and discovered a population of neutrophils that is constitutively present in the extravascular milieu. Remarkably, nasal EVN are present in large numbers in the NM independently of trauma or infection-related mechanisms, as demonstrated using imaging and germ-free mice models. While we cannot exclude the possibility that unrecognized irritants in inhaled air may contribute to neutrophil recruitment to the NM, others have recently reported that EVN are present in undisturbed tissues including skeletal muscle^62^ and lymph nodes^63^, where stimuli analogous to airborne debris are absent. Thus, we propose that neutrophil recruitment in the steady state is an integral trait of the NM, similar to other tissues.

Remarkably, our work contradicts the notion of neutrophils as a single terminally differentiated cell population^20,64^ demonstrating that nasal EVN exist as three distinct subsets. We named these subsets nN1, nN2, and nN3 and found them to be distinct at the protein and RNA levels from conventional blood and BM neutrophils, and from one another. nN1 neutrophils were found in nasal BM, a tissue that is largely ignored when studying nasal mucosa immunity. Importantly, this observation suggests previous results obtained without separating the NM from bone should be interpreted with caution, BM contamination in such preparations is likely.

Our finding that conduits connect the BM to the NM suggests a scenario where nN1 neutrophils may gain access to the NM directly from the BM, similar to skull neutrophils that migrate from skull BM to the meninges^36,37^. Thus, our study suggests that the paradigm of direct neutrophil migration from BM into target tissues without the need to enter the bloodstream is not limited to the central nervous system and can be extended to a mucosal barrier tissue. While the fate of the nN1 subset remains to be demonstrated, we postulate that this population acquires an nN2 phenotype upon arrival in the NM. In this way, the nN1 subset would provide an alternate source of neutrophils beyond the bloodstream supply. In line with this hypothesis, our pseudotime transcriptomic analysis inferred a developmental trajectory that suggests unilinear nN1 to nN2 differentiation.

Parabiosis experiments indicated that a substantial portion of nN2 and nN3 derive from neutrophils entering the tissue from blood circulation. While inflammation has been considered necessary to study neutrophil migration from blood circulation into tissues^17,65^, we revealed active neutrophil recruitment into the NM at the steady state. This recruitment is significant, as it results in neutrophils constituting approximately ∼30% of all extravascular immune cells in the NM. Notably, most studies have focused on neutrophil recruitment into inflamed tissues. The present findings thus raise questions about the molecules that are necessary for recruitment in the steady state.

Our data suggest the nN2 subset differentiates into nN3 neutrophils. The idea of neutrophil heterogeneity is relatively recent and a relevant question is whether differentiation takes place in blood or a target tissue ^64,66^. We showed that while EVN differentiation is driven by the microenvironment of the NM, as it is absent in other tissues, the process starts in the NM blood microvasculature. At the phenotypic and transcriptomic level, neutrophils in the NM microvasculature are similar to nN2 EVN, consistent with a model where differentiation starts before extravasation and continues in the extravascular space. The fact that the nN2 subset arises in the absence of inflammation raises questions about how similar these neutrophils are to their counterparts that are recruited in the context of infection or injury. Indeed, further work is necessary to determine whether nN2 neutrophils contribute to tissue homeostasis. However, we showed that nN2 neutrophils are unique in that, as opposed to the other two subsets, they cross the NM epithelium to engulf pathogens in the NC. This clearly demonstrates that nN2 are the principal first line of responders to invading pathogens.

Another surprising characteristic of nN2 is that they are not terminally differentiated. In fact, once neutrophils have access to the EV space, they appear to gradually differentiate into nN3 EVN in a process that takes place in five to six days. Of note, this indicates a neutrophil lifespan inconsistent with the notion of neutrophils being short-lived^66^. Transition is characterized by nN2 downregulation of a general neutrophil transcriptional program and upregulation of numerous gene sets associated with APC, as well as acquiring new phenotypic markers, most prominent CD11c and SiglecF.

Although SiglecF and CD11c are not markers typically associated with neutrophils, recent studies have reported SiglecF^+^ neutrophils in pathological states including cancer^67,68^ and myocardial infarction^69^. In addition, a population of SiglecF^+^ neutrophils are present in the NC in the context of allergic rhinitis^70^, *Bordetella pertussis* infection^71^, and in the uninfected neuroepithelium^72^. Consistent with these studies, we found that nN3 are highly unique to the NM. Overall, our in-depth analysis indicates that this subset consists of a unique population with a highly unusual origin, transcription programming, and associating immunologic function. Notably, we observed a pronounced acquisition of material from non-hematopoietic stromal cells by nN3, even though only limited exchange of fluorescent material between nN3 and other leukocytes was detected. Whether this reflects a preference of nN3 cells for a particular cell type or simply is a consequence of the prevailing cell types that are contacted by nN3 in the NM remains to be determined. Nonetheless, owing to the stress resulting from drastic changes in oxygenation^73^, humidity^74^, and temperature^75^, in addition to contact with airborne debris or invading pathogens, we propose that continuous tissue remodeling and repair are a hallmark of the NM. It is thus plausible that nN3 neutrophils contribute to the maintenance of homeostasis by collecting dead cell material resulting from these processes. The need for specialization to execute such functions may be a result of the high motility neutrophils possess, making the ‘cleaning’ process much more efficient than if left to macrophages or other cells with lower displacement abilities.

The ability of nN3 EVN to obtain material from the environment in combination with their phenotypic markers resembling APCs led us to propose that these cells could participate in antigen-presentation. Previous studies have described the existence of “DC-like” neutrophils that can present antigens to CD4^+^ and CD8^+^ T cells^76–79^. nN3 neutrophils, however, were able to present exogenous antigens to CD8^+^ T cells, and not CD4^+^ T cells. This is remarkable because the potential of cross-presentation by nN3 EVN could provide a new alternative for interventions in mucosal vaccinology by possibly enhancing cytotoxic T lymphocyte (CTL) induction. Importantly, our experiments showed that nN3 neutrophils induced a weaker CD8^+^ T cell proliferative response than DCs. This might be inherent to antigen-presenting neutrophils^80^. However, it is possible that a fraction of our sorted neutrophils were functionally impaired and, as a result, the number of fully functional cells was lower than in the DC control group, which had not undergone cell sorting. Alternatively, what we identify as the nN3 population might constitute more than one subset, with differing abilities to cross-present. Such considerations notwithstanding, we show that nN3s represent a cell type that cross-presents antigens exclusively to CD8 T cells.

An interesting observation is that, in addition to taking material from surrounding cells, nN3, unlike the other subsets, expresses the coinhibitory molecules PDL-1 and TIM-3 at comparable levels that those observed in DC. Furthermore, at the transcriptomic level, nN3 EVN exhibit negligible expression of IL-12, INFγ, and other cytokines essential for CTL induction. Although these properties were only assessed in the steady state, they might align with the possibility of nN3 EVN playing a modulatory role in T cell activation, with concomitant implications in immune tolerance.

As humans continue to face challenges from respiratory pathogens, our study reveals the nose as an organ possessing a complex immune arsenal prominently featuring neutrophils. These cells exhibit distinctive characteristics that set them apart from neutrophils studied in other contexts. Notably, their origins differ, with nN1 residing in BM and having access to the NM through connecting conduits. In contrast, nN2 originate from blood, to subsequently differentiate into nN3. This diversity in origin is mirrored by diversity in phenotype, evident not only in diagnostic surface proteins but also at the transcriptome level. Most importantly, functional diversity is also observed, with primarily nN2 exhibiting classical granulocyte antimicrobial activities, while nN3 neutrophils display traits that link them to adaptive immune responses. In sum, despite being part of what is traditionally considered a homogeneous cell group, in the NM EVN seem to be dramatically diverse in origin, phenotype, and function. Given the nose’s accessibility as an immune-reactive site and a growing interest in neutrophils as target cells in clinical settings^81^, our study may contribute to yet unexplored exploitation of these cells in medical interventions^82^.

## MATERIALS AND METHODS

### Mice

C57BL/6J (Jax #000664) and, β-actin RFP (Jax #005884), CD45.1 (B6.SJL-Ptprca Pepcb/BoyJ, Jax AX:002014), and BALB/cJ (Jax #000651) were purchased from Jackson Laboratories. Germ-free mice were provided by Dr. Dennis Kasper and were kept under sterile conditions before sacrificing them. Ly-6G tdTomato (Catchup) mice^24^ were kindly provided by Dr. Mattias Gunzer. LysM GFP mice^34^ were kindly provided by Dr. Thomas Graf. CD11c YFP^83^ mice were a gift from Dr. M. Nussenzweig. All experiments using mice were conducted according to national and institutional guidelines. Unless indicated, all mice were six to eight weeks old. All experiments were approved by the IACUC and COMS of Harvard Medical School.

### Nasal Cavity dissections

The nasal cavity was dissected by dislodging the nasal bone and carefully pulling it out of the skull. The skull was then pulled apart based on the sagittal axis of the head, exposing the nasal cavity. Each nasal compartment was harvested with the aid of micro tweezers and micro scissors.

### Cell preparations

Fluorescently labeled ⍺-CD45 was injected IV two minutes prior to sacrificing the mouse. Harvested nasal compartments were made into cell suspensions after digestion with collagenase D (5 mg) and DNAse (0.05 mg) in 1 mL of HBSS lysis buffer medium (1X HBSS, 10% FCS, 10 mM Hepes, 2nM CaCl_2_) incubating at 37°C for 15 min in a tube rotator. Samples were passed through a 70 µm nylon cell strainer. Cell suspensions were stained with fluorescently labeled ⍺-CD45.2 in a different color than that used for ⍺-CD45, so all leukocytes appear as CD45.2^+^ while only intravascular leukocytes appear as both, CD45.2^+^ and CD45^+^. All flow cytometry data in this study exclude doublets and dead cells.

Dead cells were identified using the LIVE/DEAD Fixable Aqua Dead Cell Stain (Invitrogen). ⍺-CD16/32 antibody was used to block Fc receptors, prior to staining with surface or intracellular markers with fluorescent antibodies mixed in FACS buffer (1% FCS, 5 mM EDTA). The Cytofix/Cytoperm Kit (BD Biosciences) a used for intracellular staining. Cells were analyzed on a Cytoflex flow cytometer (Beckman Coulter).

### Intravital Microscopy

Mice were anesthetized and placed in a stereotactic holder to immobilize the skull. The skin that covers the nasal bone was removed and a 0.5 mm burr hole was drilled through the nasal bone until the mucosa was reached. The area was monitored under a dissecting scope to ensure the tissue was not damaged (no visible hemorrhage). The area was cleaned with saline and covered with sterile GenTeal (Alcon Laboratories), and a coverslip was placed on top, along with a thermocouple to monitor local temperature, maintaining it at 37°C with a heating coil. The prepared area was placed under a 2-photon microscope to acquire multiple 3D stacks over time. Image acquisition was performed on a Pairie system at an excitation wavelength of 800 and 900 nm, from two tunable MaiTai Ti:sapphire lasers (Spectra-Physics) with physical zooming to x1-2 through an x20/0.95 numerical aperture water-immersion objective lens (Olympus). Bandpass filters of 455/70 nm, 450/80 nm, 590/50 nm, and 665/65 nm were sued to detect emitted fluorescence and second-harmonic signals. Image stacks were converted into volume-rendered three-dimensional volumes or videos using Imaris (Bitplane).

### Bulk RNAseq

Mice were dissected to obtain all regions of the nasal cavity except NALT. Cell suspensions were prepared for subsequent cell sorting in a MoFlo Astrios EQ FACS-sorter (Beckman) to isolate nN1, nN2, and nN3 neutrophils. Similarly, cell sorting was used to isolate neutrophils from blood (tail vein) and bone marrow (femur). 500-1000 cells were collected in 5 µL of TCL Buffer (Quiagen) with 1% b-mercaptoethanol. Smart-seq2 libraries were prepared as previously described^84,85^. Briefly, RNAClean XP beads (Beckman Coulter) were used to capture and purify RNA. An anchored oligo(dT) primer (5′-AAGCAGTGGTATCAACGCAGAGTACT30VN-3′) was used to select Polyadenylated mRNA and converted to cDNA through reverse transcription. With the aid of a Nextera XT DNA Library Preparation Kit (Illumina), first-strand cDNA was subjected to limited PCR amplification followed by transposon-based fragmentation. Before sequencing, amplification of samples was done through PCR for 18 cycles. Barcoded primers were used in a manner that each sample carried a specific combination of eight base Illumina P5 and P7 barcodes for amplification for 18 cycles. An Illumina NextSeq500 using 2×25 bp reads was used for paired-end sequencing.

### Bulk RNAseq analysis

The bulk RNAseq data was analyzed using an in-house bulk RNAseq data analysis pipeline. The reads were aligned to the mouse genome (GENCODE GRCm38/mm10 primary assembly and gene annotations vM20; https://www.gencodegenes.org/mouse/release_M20.html) using STAR 2.6.1d^86^.

The gene-level uniquely mapped reads quantification was calculated by featureCounts v1.6.5^87^. Samples with fewer than 7,500 genes with over ten reads were removed from the data set to eliminate the potential errors due to PCR amplification. Pairwise correlation of biological samples was performed to identify poor-quality samples; any replicates that fell below 0.85 cutoff were discarded. Data from all samples was combined into a raw reads count table for differential gene expression using DESeq2.

Genes whose total read count summed over all the samples fell below 50, were trimmed from the table as noise. The raw reads count tables were normalized by DESeq2’s “median of ratios” method. Each population was pairwise contrasted against each other with 0.05 FDR. Genes with an expression fold-change >2 and Benjamini-Hochberg adjusted p-value < 0.05 were considered significant and used in further analysis.

### Single Cell RNAseq

Naïve LysM GFP mice were euthanized 3 minutes after IV injection of an oligo-barcoded (TotalSeqB) anti-CD45 antibody (PN). After harvesting nasal tissue and processing as described above, cells were stained with oligo-barcoded antibodies against Ly6G (PN), CD11b (PN), CD11c (PN), and Siglec-F (PN) and sorted to collect GFP^bright^ cells. 20,000 oligo-barcoded antibody-stained GFP^bright^ cells were then processed for single-cell RNA-sequencing using 10x Genomics Chromium Next GEM Single Cell 3’ Kit v3.1 and Feature Barcoding Kit with dual indices per the manufacturer’s instructions. For sequencing, the antibody barcode library was pooled at a ratio of 1:5 with the gene expression library and sequenced on a NextSeq 2000 (Illumina) to an average depth of 34,000 reads per cell.

### Single Cell RNAseq analysis

Sequencing data was processed using the 10x Genomics cellranger pipeline (v7.0.0) via the Cumulus platform^88^ to generate count and barcode matrices. Subsequent analysis was performed using Seurat (v5.0.1) in R. In brief, following filtration of cells with fewer than 500 UMIs or 300 genes, cells were processed via scTransform, PCA, clustering, and UMAP embedding to reveal 7 clusters across two identified cell types by gene expression: neutrophils and macrophages. All macrophage clusters were then removed from the subsequent analysis, and the neutrophils were reprocessed through clustering and UMAP embedding. Following the removal of two multiplet clusters expressing monocyte (*Ms4a6c*) or odorant-binding protein genes, the remaining neutrophils were reprocessed a final time to reveal 7 clusters. Each cluster was then evaluated on the expression of *Ly6g*, *Itgam, Itgax,* and *Siglecf* and matching protein expression (via barcode expression matrix) to assign identity to either the nN1, nN2, or nN3 subset. Pseudotime analysis of nasal mucosa neutrophils was performed using Monocle 3^48,89,90^. Seurat-processed data, specifically the normalized gene expression matrix and UMAP embeddings, were imported into Monocle3. A principal graph was fitted to the data to learn the trajectory of transcriptional changes. The cells were then ordered in pseudotime by assigning the beginning of the trajectory to a root node in the nN1 clusters.

### Gene set enrichment analysis for N3 and Immgen datasets

To compare the differentially expressed genes (DEGs) of the N3 population to other immune populations, we downloaded raw count data of several immune cell types from Immgen ^53^. Differential gene expression analysis was performed using the DEseq2 pipeline by contrasting each cell type against all others. The resulting DEGs were then sorted by log-fold change and used as ranks for pre-ranked fast gene set enrichment analysis using the “fgsea” R-package^91^. The DEGs of the N3 population (see above) were then used to calculate normalized enrichment scores for each of the pre-ranked pathways.

### Parabiosis

Anesthetized 8-week-old congenic CD45.1 and CD45.2 mice (age- and sex-matched) were shaved on the site of the incision. Matching incisions were made on the skin from the olecranon to the knee joint of each mouse. A suture was used to join the olecranona and knees from both partners. Staples (Stoelting) were used to approximate the dorsal and ventral skins of both mice. Animals remained conjoined for five weeks. Samples were harvested after this time.

### Hematopoietic progenitors

The mucosa from the aNR, pNR, NT, and SE regions (not shown) were pooled together. The ET, was harvested separately and further processed to separate mucosa from bone. Cell suspensions were either stained for hematopoietic progenitor markers for flow cytometry or used in colony-forming unit (CFU) assays. The MT region was excluded to avoid the risk of bone from this region contaminating cell suspensions. For CFU assays, cell suspensions from fluorescently labeled mice were mixed with methylcellulose-containing MethoCult M3534 (Stemcell Technologies) in various dilutions and incubated at 37°C. Cell fluorescence was used to differentiate CFU from debris. Colonies were counted under a fluorescence dissecting microscope and passed through a 70 µm filter for FACS analysis to determine cell size through forward light scattering.

### Pathogens

#### Construction and growth of *Chlamydia muridarum*-dsRed

A DNA fragment corresponding to the *dsRed* open reading frame (ORF) was amplified by PCR from pGEN-Pem7-DsRedT3mut using primers GroESL DsRed START 5 (5’→3’, GAATATAAAAATACGAGGAGCTTAAACATGGCGAGCAGTGAGGACATC)and GroESL DsRed STOP 3 (5’→3’, CTATTTGTTCCCATTAGAGGAACTAAAGGAACAGATGGTGGCG). DNA fragments corresponding to the intergenic regions upstream (Gro Promoter) and downstream (Gro Terminator) of the *groESL* operon were amplified by PCR from *C. trachomatis* LGV L2 genomic DNA by using primers GroESLProm Age 5 plus GroESL DsRed START 3 (5’→3’, GATGTCCTCACTGCTCGCCATGTTTAAGCTCCTCGTATTTTTATATTC (Gro Promoter)) and GroESL DsRed STOP 5 (5’→3’, CGCCACCATCTGTTCCTTTAGTTCCTCTAATGGGAACAAATAG) plus GroESLTerm Age 3 (Gro Terminator), respectively. A DNA fragment corresponding to Gro Promoter-dsRed-Gro Terminator was then amplified by overlapping PCR using the primers GroESLProm Age 5 (5’→3’, ACCACCGGTATTTTTAAAAATAGCAGTTGATCATGCC) and GroESLTerm Age 3 (5’→3’, ACCACCGGTAGAAAAGGATGGTCGTAAGCACTAG) and cloned into the AgeI site of p2TK2_Spec_-Nigg ^92^. *C. muridarum*-dsRed transformation and plaque purification was then performed as described previously^92^. *C. muridarum*-dsRed was propagated in McCoy cells using 500 µg/ml spectinomycin and elementary bodies (EBs) isolated as previously described^93–95^. Aliquots of purified EBs were stored at - 80°C in SPG buffer containing 250 mM sucrose, 10 mM sodium phosphate, and 5mM L-glutamic acid and thawed immediately prior to use.

#### Construction of Streptococcus pneumoniae GFP

Pneumococci were grown in Todd-Hewitt broth supplemented with 0.5% yeast extract (THY) or on plates with tryptic soy agar with 5% sheep blood (TSA). Antibiotics (300 µg/ml of kanamycin, 600 µg/ml of streptomycin, and 4 µg/ml of chloramphenicol) were added as needed. Every pneumococcal mutant was derived from a spontaneous streptomycin-resistant (Smr) strain (T4) derived from serotype 4 clinical isolate TIGR4. DNA cloning strategies were based on overlapping PCR and other previously described techniques^96^. The hlpA-GFP strain was generated using the bicistronic Janus cassette in a two-step method^97^. The ORF of gene *SP_1113* was initially replaced with the Janus cassette then the fusion of hlpA-sfGFP provided to us by the Veening team was used to replace the Janus cassette. Fluorescent bacteria were screened using a dissecting scope equipped with excitation to detect GFP in colonies.

#### Inoculations

Mice were inoculated with *C. albicans* (10^6^ CFU), *C. muridarum* (10^6^ IFU), or *S. pneumoniae* (10^6^ CFU) using the intranasal route. Pathogens were instilled into the nares (5 µL per nare) of partially anesthetized mice.

### In vitro T cell expansion

DC were used as control and were generated by Flt3L-induction of bone marrow-derived DC as previously described^98^. Briefly, cells from BM (femora and tibiae) obtained from C57BL/6J mice were plated with 100 ng/ml or recombinant Flt3L (R&D systems) in RPMI1640 medium (Gibco) with 10% FBS (Gemini Bio), β-mercaptoethanol, glutamine, and penicillin–streptomycin. DCs were harvested and used on days 9-10. CD4+ and CD8+ T cells were obtained from cell suspensions made from mouse spleens that were further purified using the CD4+ T Cell Isolation Kit (Milteny Biotec), and CD4+ T Cell Isolation Kit (Milteny Biotec).

Sorted nN1, nN2, nN3, blood neutrophils, and DC were pulsed with OVA (10 µg/mL) for 3 hours, washed, and placed in a 96 well-plate (2500 cells/well). These cells were cultured with purified OVA-specific CD4+ or CD8+ T cells that were stained with a cell proliferation dye (Cell Trace Violet, ThermoFisher Scientific) for three days (100,000 cells/well). Proliferation was assessed through FACS.

