## Supplementary figures, figure legends, and tables for "Constitutive immune surveillance of nasal mucosa by three neutrophil subsets with distinct origin, phenotype, and function"

SUPPLMEMENTARY FIGURES

#### SUPPLEMENTARY FIGURE 1

A

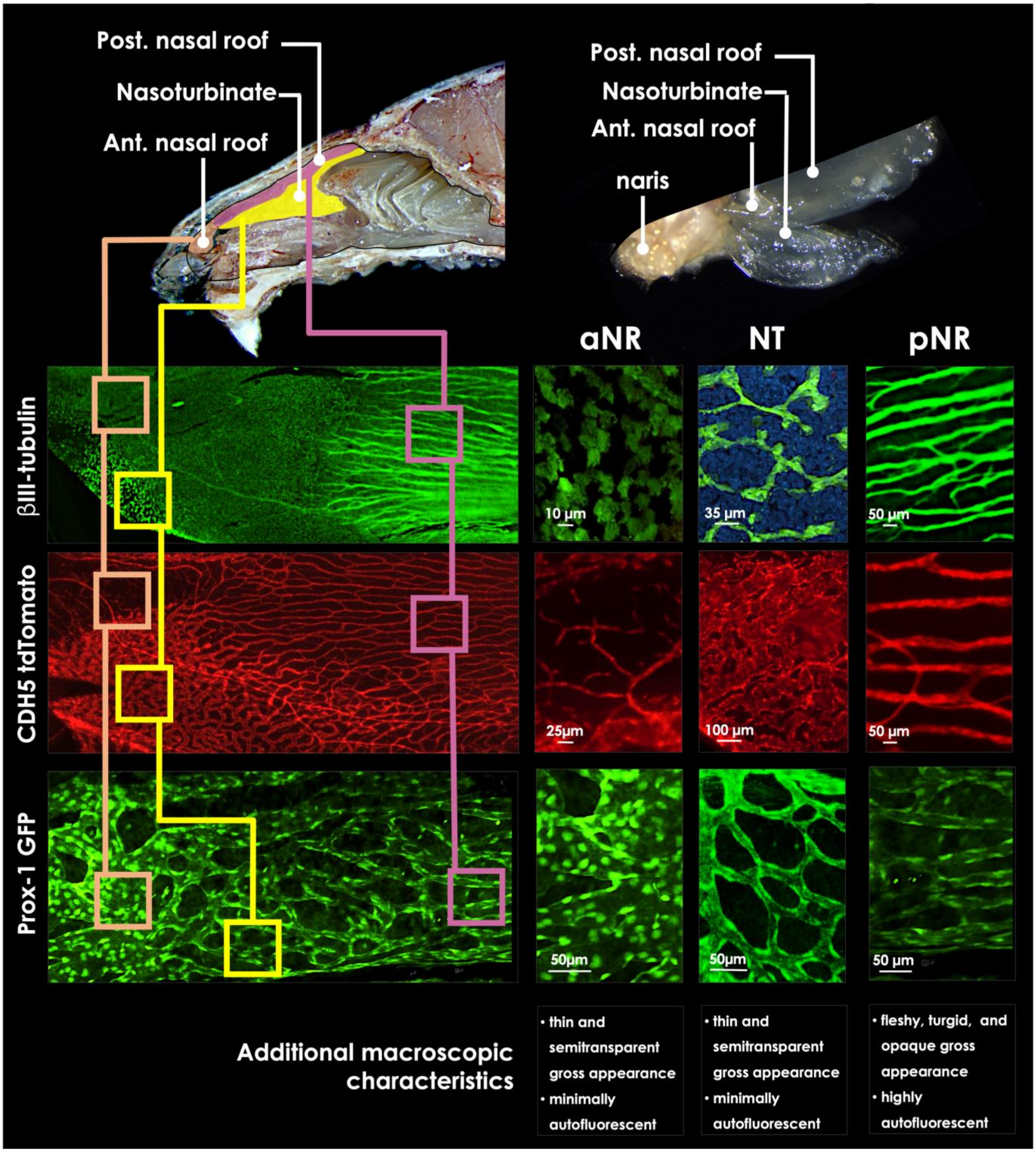

B

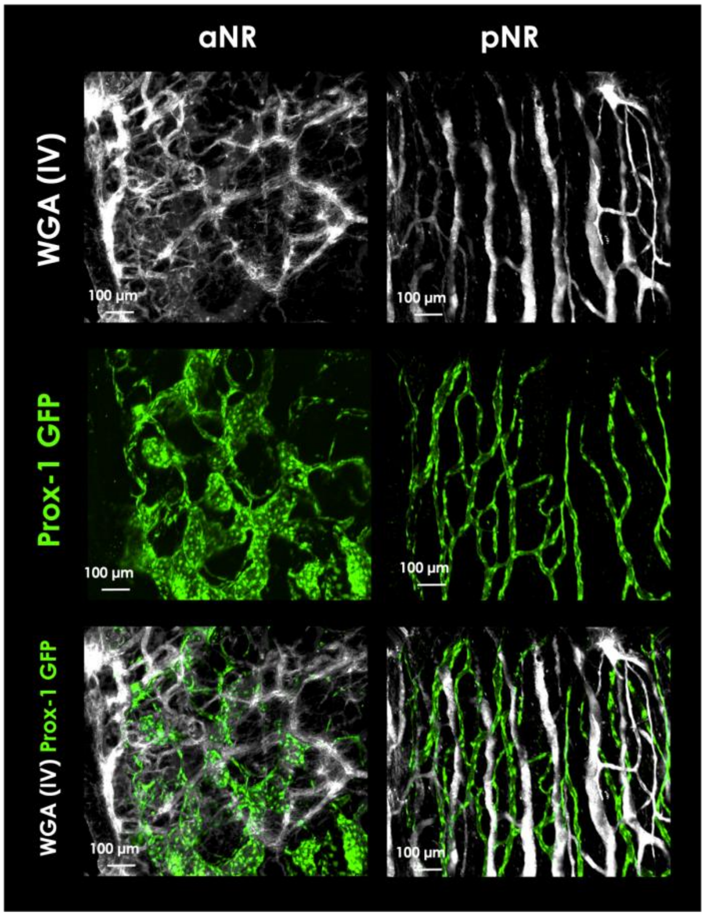


**Supplementary Figure 1. The mucosa of the aNR, NT, and pNR regions can be identified by different parameters. (A)** The upper left side of the panel shows the location of these regions in the NC. The upper right side of the panel displays an image of the three regions harvested from a mouse. The three regions conform to a single layer of tissue. The apparent separation between the pNR and the NT mucosa is due to unavoidable disruption of the tissue when harvesting it from the NC. The subpanels of the left display a fraction of the tissue where the three regions converge, with squares of different colors indicating the approximate location of each region for each image. The subpanels on the right display images from each region. Nerve bundles and cell cilia, both identified by βIII-tubulin staining (green), were one of the criteria to establish differences between the three regions. Prominently large nerve bundles of approximately 50-75 µm in caliber, running along the length of the nasal bone and parallel to one another, were visualized in the pNR mucosa. No ciliated cells were observed in this region. Ciliated cells forming a patchy pattern and very dim signal (from both, neurons and ciliated cells) were observed in the aNR region. No nerve bundles were observed in the mucosa of the NT region. However, signal from ciliated cells was strong and well-defined. Ciliated cells formed conglomerates suggesting they are not evenly distributed in the NT mucosa. Blood vasculature (CDH5 tdTomato, red) in the aNR region appears as branching vessels of approximately 10 µm in caliber. The NT mucosa contained thick meandering vessels of approximately 100 µm in caliber that resembled sinusoid vessels. In the pNR mucosa vessels of approximately 50 µm in caliber run parallel to one another and along the length of the nasal bone, with unidirectional blood flow in a caudal to rostral direction, moving towards the nostrils (determined by observation of a live mouse under the microscope, not shown). In the aNR mucosa, lymphatic vessels (Prox-1 GFP) were prominently large (100-150 µm in caliber) and prominently bright in the Prox-1 GFP mouse. Lymphatic vessels in the NT, with a caliber of approximately 20-30 µm, appeared in a reticular arrangement. The lymphatic vasculature in the pNR mucosa consisted of semi-parallel vessels of approximately 25 µm in caliber that appeared dim in the Prox-1 GFP mouse. Additional characteristics described at the end of the panel were determined by observation under dissecting and fluorescence microscope. **(B)** Images from the aNR and pNR showing blood vasculature (after IV injection of labeled WGA, minutes before sacrificing the mouse) and lymphatic vasculature (Prox-1 GFP), simultaneously.

### SUPPLEMENTARY FIGURE 2

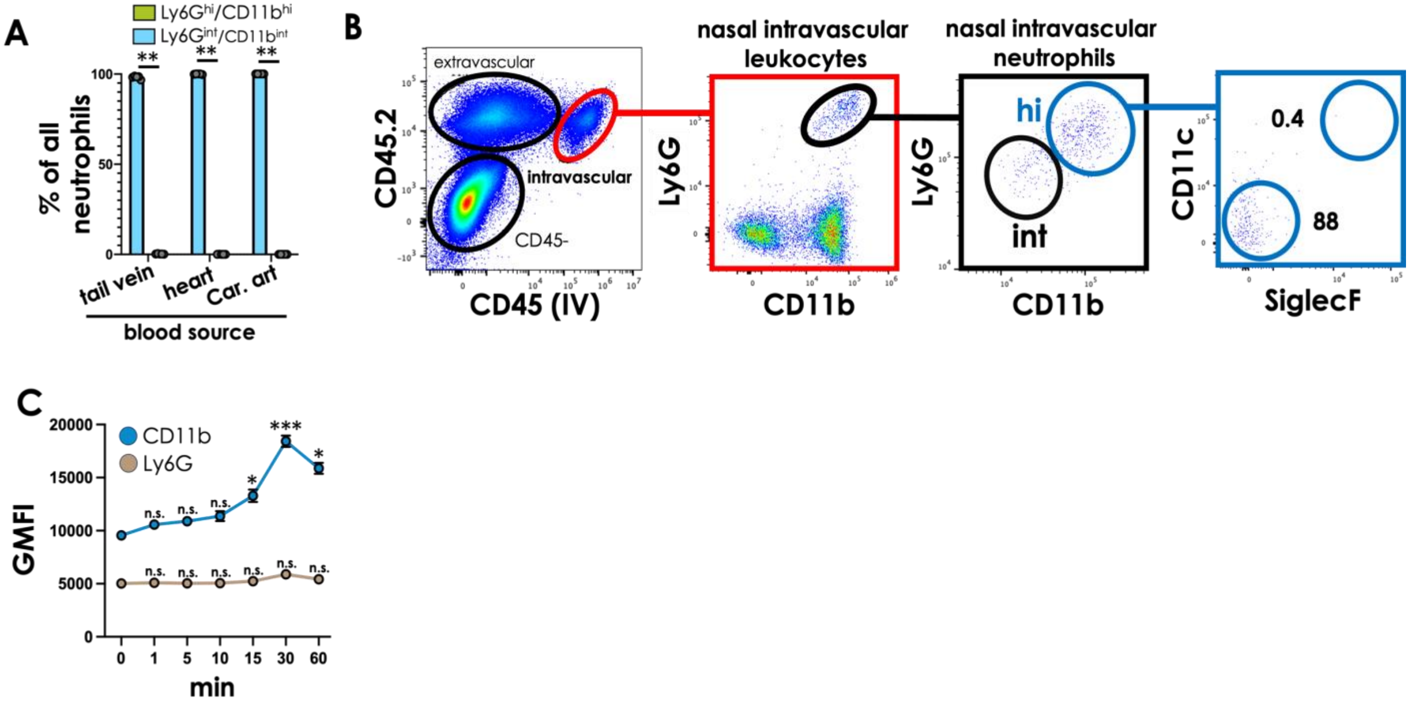


**Supplementary Figure 2. Nasal intravascular neutrophils are CD11c$^-$SiglecF$^-$. (A)** Frequency of CD11b$^{hi}$Ly6G$^{hi}$ and CD11b$^{int}$Ly6G$^{int}$ neutrophils from blood obtained from the tail vein, the heart, or the carotid artery (Car. art, which supplies blood to the nasal mucosa). All neutrophils from blood, regardless of source, exhibit a CD11b$^{int}$Ly6G$^{int}$ phenotype. Mann-Whitney test. Differences between groups are considered significant when p<0.05. **P<0.01,. Circles represent a single mouse; bars represent the mean of the group and error bars represent SEM. **(B)** Representative dot plot depicting SiglecF and CD11c expression of CD11b$^{hi}$Ly6G$^{hi}$ nasal IVN. **(C)** Geometric mean fluorescence intensity (GMFI) of CD11b (blue) and Ly6G (brown) staining on BM neutrophils at different time points after treatment with LPS in vitro. One out of two independent experiments, with three wells per time point. Circles represent the average value of the three wells. Differences between zero time points and other time points were determined with a two-way ANOVA test and were considered significant when p<0.05. *P<0.01, ***P<0.001.

#### SUPPLEMENTARY FIGURE 3

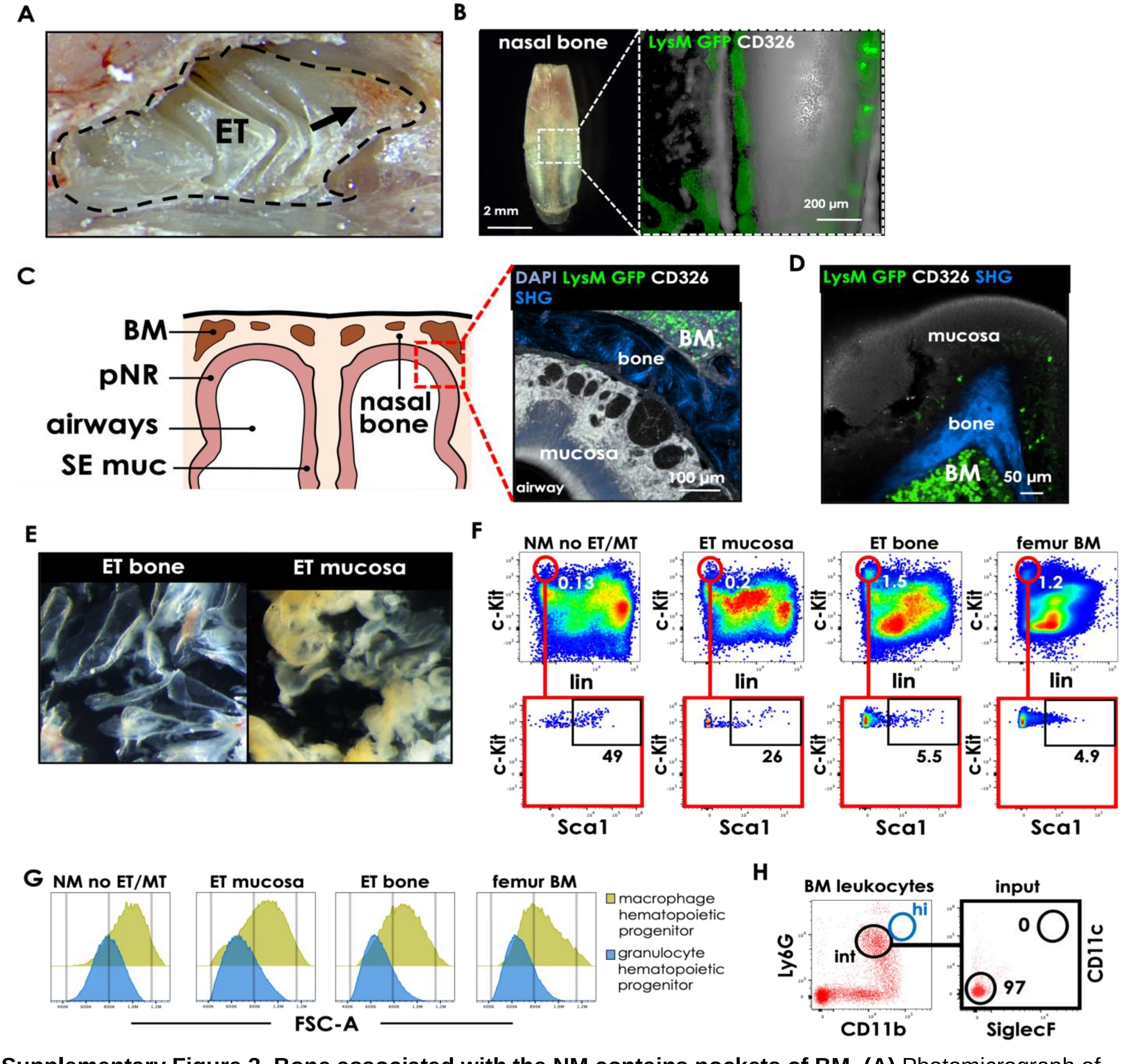


**Supplementary Figure 3. Bone associated with the NM contains pockets of BM. (A)** Photomicrograph of the ET, a nasal region with zones that are reminiscent of BM (arrow). **(B)** Photomicrograph of the nasal bone with zones (upper half) that are reminiscent of BM (left side of the panel). The right side of the panel shows a maximum projection of cleared tissue corresponding to the nasal bone of a LysM GFP mouse imaged through light sheet microscopy. The dotted rectangle on the left indicates the area of the nasal bone that is imaged on the right side. Pockets of BM are visualized in green (through the fluorescence of GFP$^{bright}$ neutrophils). The epithelium and some autofluorescence from bone synarthroses are visualized in white. **(C)** Left, schematic depicting a frontal section of the NC, with BM embedded in the nasal bone, above the mucosa of the pNR region. Right, photomicrograph of a cross-section from an area corresponding to the dotted red line in the schematic (left) of a LysM GFP mouse. BM appears in the upper right corner, visualized through the fluorescence of $GFP^{bright}$ neutrophils. BM is adjacent to the pNR mucosa with bone separating the two compartments. Bone appears in bright blue. Epithelium and nuclei appear in white and blue, respectively. **(D)** Photomicrograph of a cross-section of the ET region from a LysM GFP mouse. As in the nasal bone, BM is also present in the ET and is visualized through the fluorescence of $GFP^{bright}$ neutrophils. BM is separated from the ET mucosa by bone. Epithelium and bone appear in white and blue, respectively. For **C** and **D**, bone is visualized through second harmonics generation (SHG) using 2-photon microscopy. **(E)** Photomicrographs depicting ET bone (left) and ET mucosa (right) after separation of the two tissues. The mucosa was gently pulled apart from bone with a fine micro brush and the action of PBS at high pressure. The bone fraction contains visible red areas corresponding to BM. **(F)** Representative dot plots displaying $lineage^-$ (i.e.: $CD3^-$, $CD11b^-$, $CD11c^-$, $CD19^-$, $Ly6G^-$, $NK\text{-}1.1^-$, $NKp46^-$, and $Ter\text{-}119^-$), $c\text{-}Kit^+$ cells (upper row) that are also $Sca1^+$ (lower row). Gated on live cells (determined by live/dead staining) that were $CD45^+$. **(G)** Representative histograms displaying forward scatter of cells from colonies harvested after cell suspensions from aNR, pNR, NT, and SE region mucosa (pooled together), ET mucosa, ET bone, and femur BM were incubated in semisolid medium for the expansion of granulocyte-macrophage progenitors. Both, granulocyte (smaller size, blue) and macrophage (larger size, green) hematopoietic progenitors are present. Gated on RFP+, as tissues were obtained from $\beta$-act RFP mice to easily distinguish colonies from debris. **(H)** Representative dot plot of BM leukocytes. Adoptively transferred neutrophils obtained from BM (input) are $CD11b^{int}Ly6G^{int}$ and $CD11c^-$ $SiglecF^-$, to neutrophils from blood.

#### SUPPLEMENTARY FIGURE 4

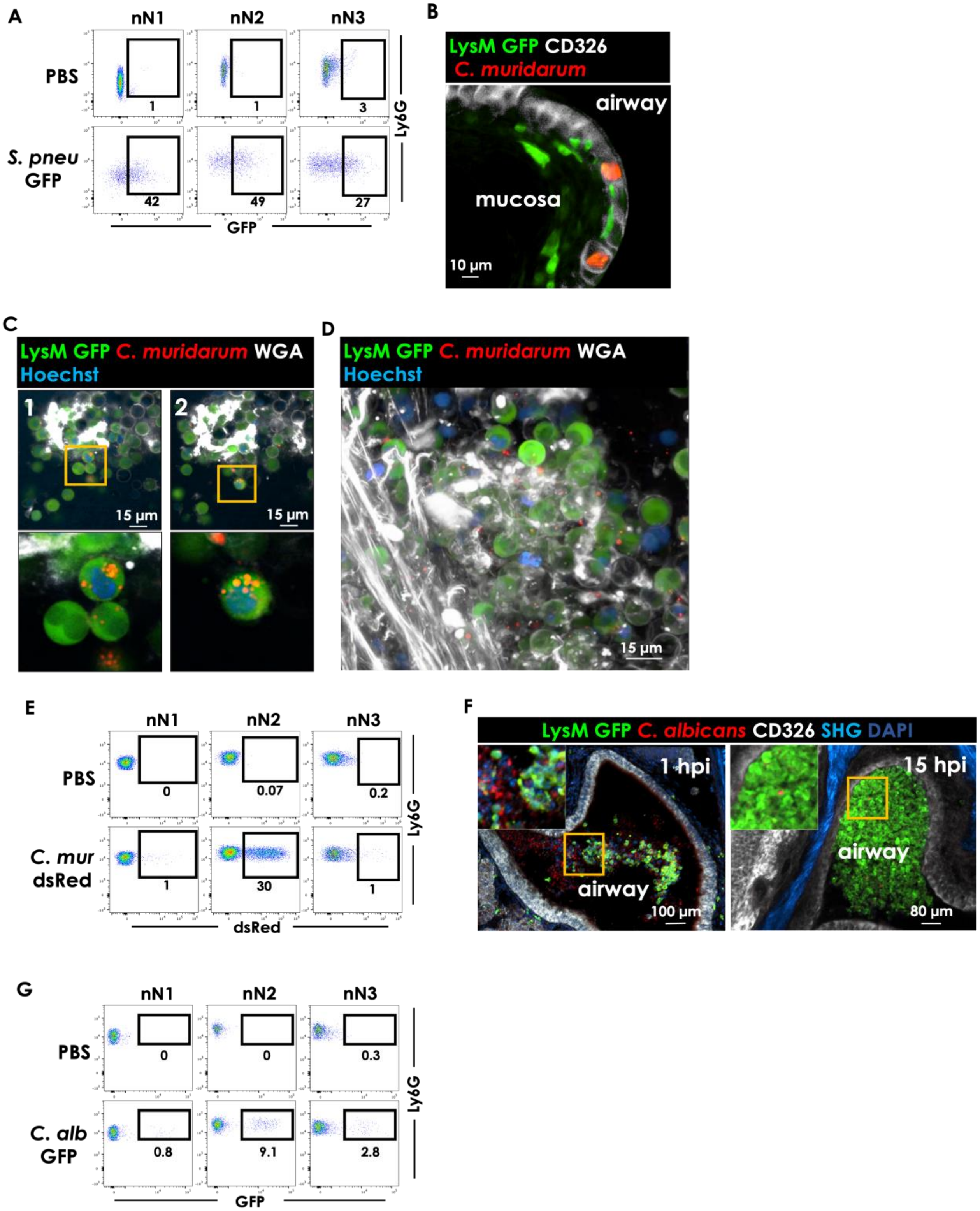


**Supplementary Figure 4. nN2 neutrophils associate with different pathogens. (A)** Representative dot plots of sorted nN1, nN2, and nN3 neutrophils after *in vitro* incubation with PBS or *S. pneumoniae* GFP (multiplicity of infection 0.1) for 3 hours. **(B)** Representative photomicrograph of the NM from a LysM GFP mouse that was inoculated intranasally with *C. muridarum* dsRed (red) at 24 hpi. Neutrophils appear in green, the epithelium (CD326) in white, and *C. muridarum*, forming inclusion bodies, in red. **(C)** Representative photomicrographs of two consecutive optical slices (1 and 2) from a stack obtained similar to that shown in Supplementary video 2, where a LysM GFP mouse was inoculated intranasally with *C. muridarum* dsRed (red). At 24 hpi, nuclear stain (Hoechst, blue) was injected intravenously and 5 µL of WGA (white) were instilled into the nostrils, to visualize mucus. The mouse was sacrificed 5 minutes after WGA and Hoechst treatment, the NC was dissected and contents of the lumen (i.e., airways) were imaged in situ using 2-photon microscopy. The orange square demarcates the area magnified below each image. *C. muridarum* appears inside neutrophils (green). **(D)** Representative photomicrographs of a maximum projection from Supplementary video 2. Experimental conditions were the same described in C. The parallel elongated structures on the left are mucus attached to the epithelium. The majority of neutrophils are suspended in mucus of the airways. **(E)** Representative dot plots of nN1, nN2, and nN3 neutrophils 48 h after IN instillation of PBS or *C. muridarum* dsRed. **(F)** Representative photomicrographs of the NC from LysM GFP mice inoculated intranasally with *C. albicans* tdTomato (red). The tissues were harvested at one and 15 hpi. Neutrophils appear in green, the epithelium (CD326) in white, nuclei in blue (DAPI) and bone (SHG) in bright blue. **(G)** Representative dot plots of nN1, nN2, and nN3 neutrophils 15 h after IN instillation of PBS or *C. albicans* GFP. For A, D, and F, gates were drawn using the PBS groups as a reference to avoid confounding effects due to autofluorescence.

### SUPPLEMENTARY FIGURE 5

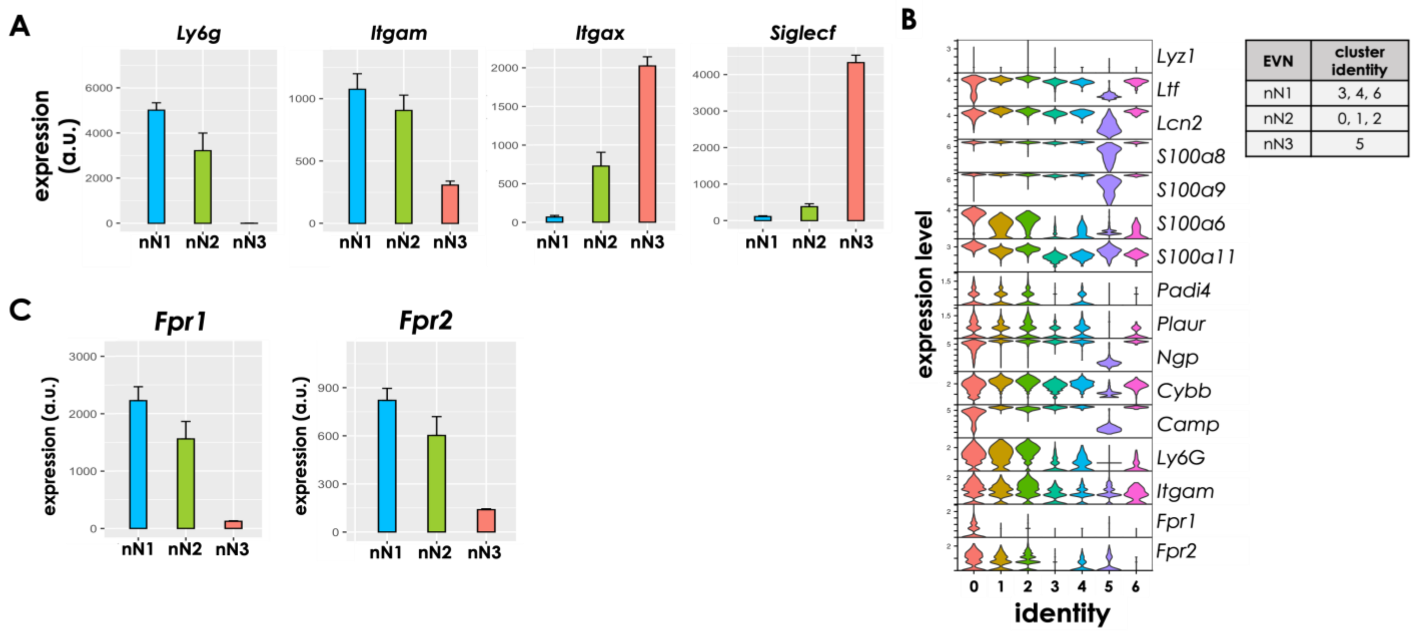


**Supplementary Figure 5. Transcriptome analysis of NM neutrophil subsets.** (**A**) Expression of *Ly6g*, *Itgam*, *Itgax*, and *Siglecf* in nN1, nN2, and nN3 EVN as determined by bulk RNAseq (see Fig. 6B). (**B**) Violin plot indicating the expression of 15 selected genes that are commonly ascribed to neutrophil functions (see Fig. 6G) and the genes *Fpr1* and *Fpr2*, involved in neutrophil detection of microbial pathogens (see Fig. 6H), as determined by scRNA seq. The table on the right indicates the cell clusters that were ascribed to the three EVN subsets. Cluster identity based on UMAP shown in Fig. 6C. No expression of the other genes involved in neutrophil detection of microbial pathogens shown in Fig. 6H was detected by scRNAseq. (**C**) Expression of *Fpr1* and *Fpr2* as determined by bulk RNAseq. For **A** and **C**, each column represents the mean expression of four independent biological replicates of sorted nN2, nN2, and nN3 EVN.

### SUPPLEMENTARY FIGURE 6

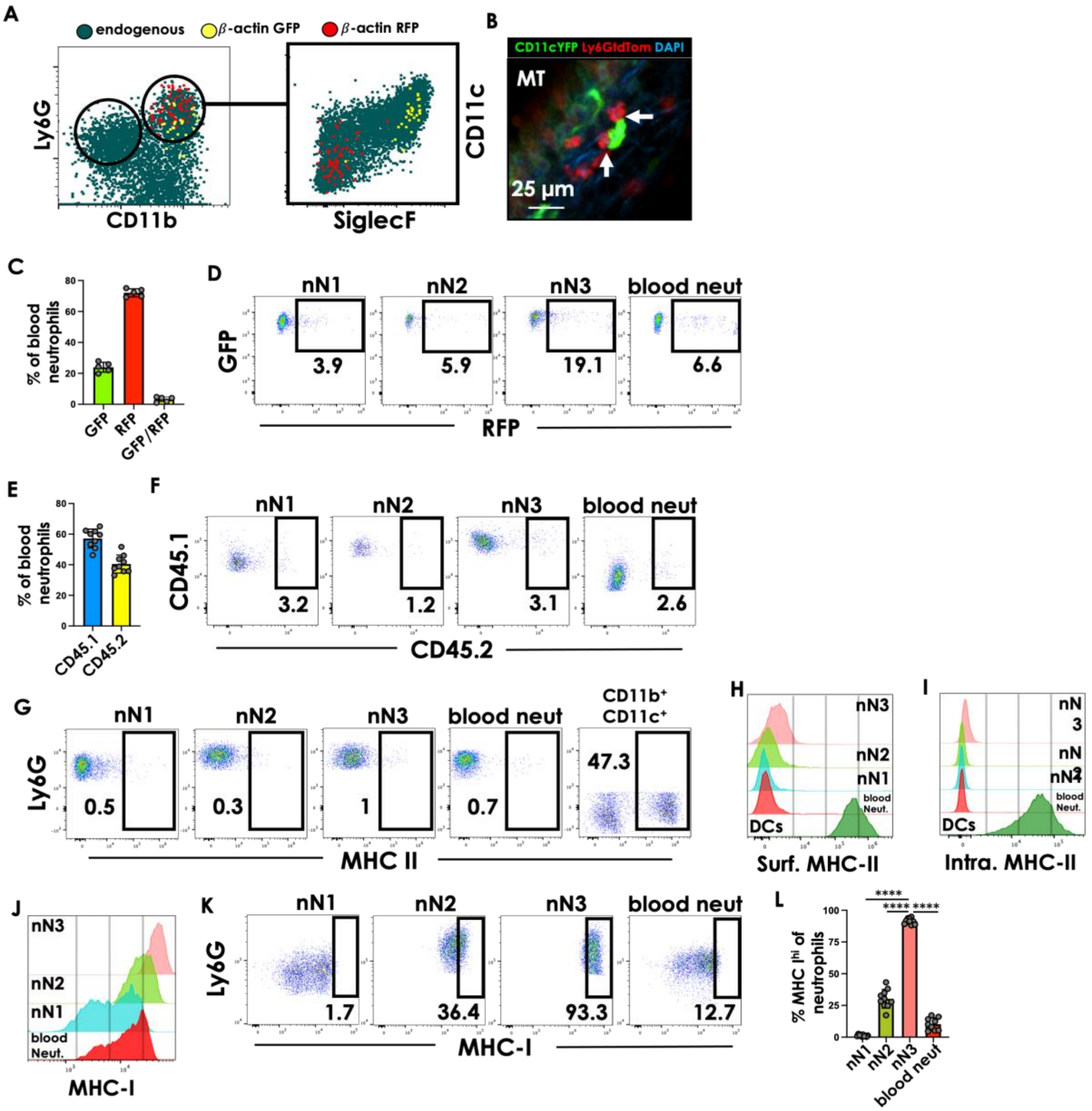

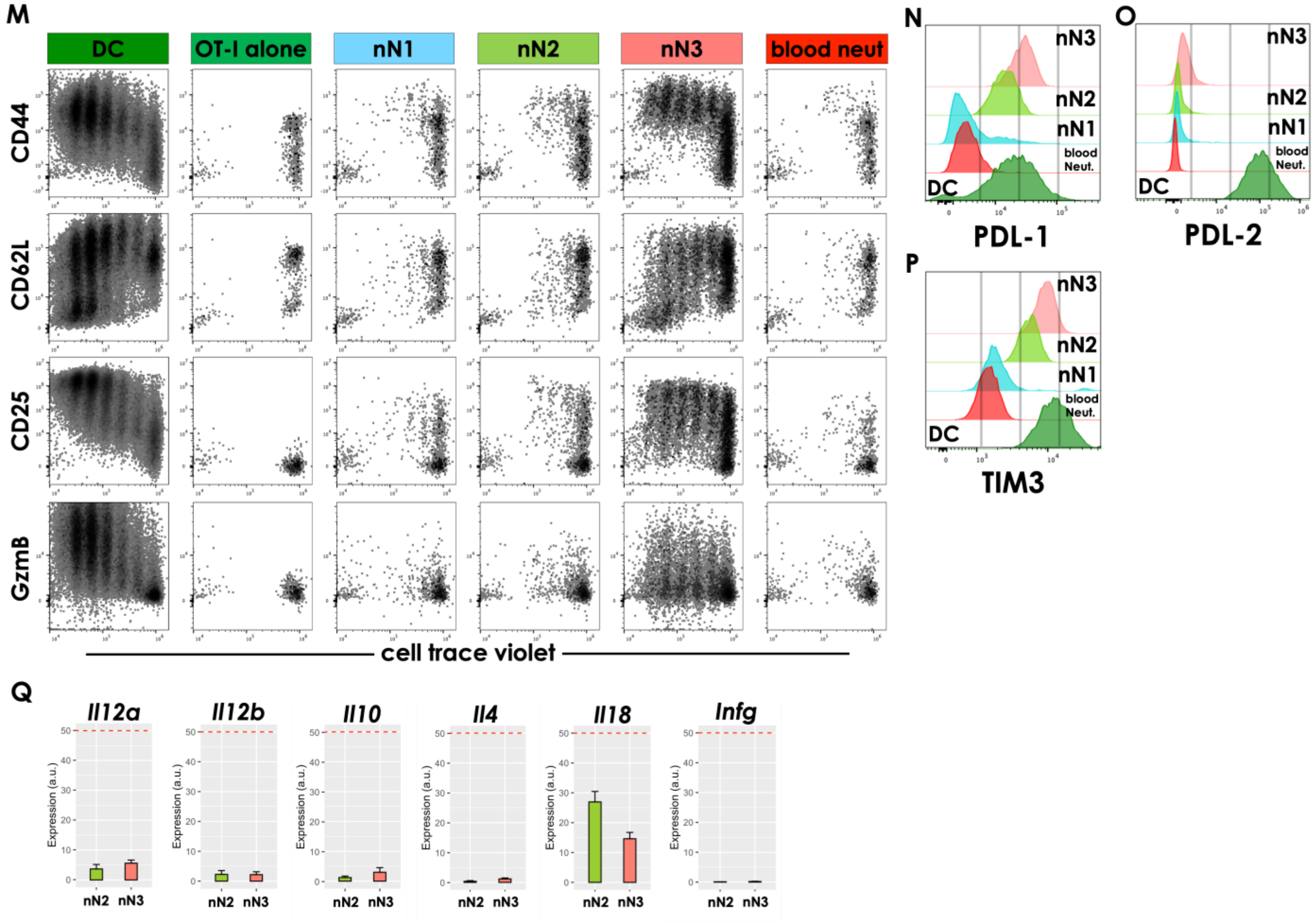


**Supplementary Figure 6. nN3 EVN assume phenotypic and functional features of APCs**

**(A)** Sorted BM neutrophils from β-actin GFP and β-actin RFP mice were injected IV into a non-fluorescent mouse at day zero and day four, respectively. NM cell suspensions were made on day six. At this time point, β-actin GFP (yellow dots) and β-actin RFP (red dots) neutrophils exclusively exhibited an nN3 and nN2 phenotype, respectively. Endogenous neutrophils appear in dark green. **(B)** Representative photomicrograph showing close interactions between Ly6G tdTomato neutrophils (red) and CD11c YFP cells (green) in the NM. Points of apparent contact between the two cells are marked with arrows. The image corresponds to a single optical section from a fixed whole mount tissue. **(C)** Frequency of β-actin GFP and β-actin RFP neutrophils in blood from mixed BM chimeric mice. β-actin GFP and β-actin RFP BM was mixed and transplanted into non-fluorescent lethally irradiated recipient mice. **(D)** Representative dot plot of β-actin GFP nN1, nN2, nN3, and blood neutrophils from the mixed BM chimera mice described in C **.** Gating on $GFP^+$ $Ly6G^+$ $CD11b^+$ cells **(E)** Frequency of CD45.1 and CD45.2 neutrophils in blood obtained from mixed BM chimeric mice where BM from congenic CD45.1 and CD45.2 mice was mixed and transplanted into lethally irradiated CD45.2 mice. **(F)** Representative dot plot of nN1, nN2, nN3, and blood neutrophils from the mixed BM chimera mice described in E. Gating on $CD45.1^+$ $Ly6G^+$ $CD11b^+$ cells**. (G)** Representative dot plot displaying expression of MHC-II in nN1, nN2, nN3, and blood neutrophils. $CD11c^+$ $CD11b^+$ $Ly6G^-$ cells (likely DCs) used as positive control. (**H-I**) representative histogram showing nN1, nN2, nN3, and blood neutrophils expression of surface (**H**) and intracellular (**I**) MHC-II. DCs from the NM used as reference. **(J)** Representative histogram showing nN1, nN2, nN3, and blood neutrophil expression of MHC-I. **(K)** Representative dot plot showing expression of MHC-I in nN1, nN2, nN3, and blood neutrophils. All cells express MHC-I but only nN3 neutrophils express it at high levels. Gates were drawn using blood neutrophils as a reference. Cells inside the drawn gate were identified as "hi" for MHC-I. **(L)** Quantification of K, displaying the frequency of MHC-I$^{hi}$ for nN1, nN2, nN3, and blood neutrophils. One-way ANOVA with differences between nN3 and the other groups considered significant when p<0.05. ****P<0.0001. **(M)** Representative dot plots of cell trace violet-labeled OVA-specific OT-I CD8 T cells that were incubated for three days with OVA-pulsed sorted nN1, nN2, nN3, blood neutrophils, or Flt3L-induced BM-DCs. For C, E, and L, circles represent a single mouse; bars represent the mean of the group, and error bars represent SEM. Data from one representative experiment shown for C and two pooled experiments for E and L. For A, D, F, G, K, and M, gates were drawn on live (based on live/dead stain) neutrophils ($CD11b^{+}Ly6G^{+}$), excluding doublets. (**N-O**) Representative histogram showing nN1, nN2, nN3, blood neutrophil, and DC expression of PDL-1 (**N**), PDL-2 (**O**), and TIM3 (**P**). (**Q**) Expression of *Il2a*, *Il2b*, *Il10*, *Il14*, *Il18,* and *Infg* in nN2 and nN3 EVN as determined by bulk RNAseq.

#### SUPPLEMENTARY TABLES

**Supplementary Table 1.** The most relevant genes driving variance in PC1 and PC2, in the PCA analysis shown in Fig. 3D.

| PC | % variance explained | Top 100 Genes separating samples |
|---|---|---|
| **1** | **56.43** | Cyp2a5, Omp, Gm14744, Obp1a, Cyp2f2, 5430402E10Rik, Obp1b, Sec14l3, Stoml3, Cyp2g1, Gm14743, Bpifb3, Scgb1c1, Lcn11, Egr1, Mup4, S100a5, Map1b, Bpifa1, Obp2a, Cbr2, Chga, Gm14750, Umodl1, BC051076, Gfy, Ccrl2, Wfdc18, Rgs5, Car6, Muc2, Cyp2a4, Vmo1, Krt18, Epas1, Cxcl2, Obp2b, Cnga2, Nsg1, Scgb2b27, Ckb, Plekhb1, Fstl5, Sparc, Fcrls, Gpx6, Pcp4l1, Bpifb4, Mup5, Ppp1r15a, Ak1, Rtn1, Ptgs2, Rgs1, Rtp1, Atp1b1, Aox2, Tspan7, Cdkn1a, Epcam, Kirrel2, Tubb3, A2ml1, Ptprs,Bpifb6, Ebf2, Cyp2a21−ps, Aplp1, Cyp1a2, Hist2h2be, Gas6, Glb1l2, Scgb1b27,Col1a1, Col1a2, Kif1a, Reg3g, Flrt1, Padi4, Ckmt1, Npdc1,Tmem176b, Ctxn3, Gng13, Plxnb2, Cldn5, C1qb, Stmn2, Sparcl1, Jun, Ebf4, Cx3cr1, Ece1, Sell, Adamdec1, Ly6g, Cnga4, Cldn3, Pon1, Aqp3 |
| **2** | **18.17** | Camp, Ngp, I830127L07Rik, Ltf, Chil3, Ly6g, 4930438A08Rik, Adpgk, Itgb2l, Ifitm6, Cd177, Serpinb1a, Olfm4, Mmp8, Gm6522, Ly6c2, Chil4, F730016J06Rik, Mgst2, Acvrl1, Inhba, 1700047M11Rik, Thbs1, Cybb, Ear2, Chil5, Olfml2b, Hba−a2, Lbp, Cldn15, Hbb−bt, Padi4, Igkc, Fcnb, Hbb−bs, Lcn2, Gm14744, Chil1, Top2a, Ccna2, Ly6g5b, Ceacam1, Anxa1, St3gal5, Cdc20, Ube2c, 5430402E10Rik, Tinagl1, Cd74, Hba−a1, Gm33326, H2−Ab1, Pnpla1, Mlst8, Itgax, C3, Reck, Wfdc21, Zmpste24, Siglecf, Serpinb10, H2−Aa, Tmem40, Ptgir, Osm, Rasl11b, Syne1, Ccnb1, Capg, Anxa3, Lta4h, Ldlr, Ccnb2, Slc17a9, Dstn, Mmp25, Flot2, Ceacam10, Krt83, Obp1a, Slc31a2, Ano10, Pxylp1, Obp1b, Rrm2, Cdca8, Lcn11, H2−Eb1, Mogat2, Gm14750, S1pr4, Trp53inp2, Socs3, Prom1, Ankrd22, BC051076, Ltb4r1, Abca13, Alas2, Retnlg |

**Supplementary Table 2.** Most relevant enriched gene ontology (GO) terms from genes that are downregulated in nN3 respective to nN2 neutrophils. Terms are organized by fold enrichment, from highest to lowest. Only the 15 biological process GO terms with highest positive fold enrichment are shown. Raw P value is shown for each term.

| # | GO biological process (downregulated in N3) | +/- | Fold Enrichment | raw P value |
|---|---|---|---|---|
| 1 | positive regulation of neutrophil activation | + | 21.87 | 3.11E-06 |
| 2 | cellular response to diacyl bacterial lipopeptide | + | 21.87 | 1.15E-03 |
| 3 | response to diacyl bacterial lipopeptide | + | 21.87 | 1.15E-03 |
| 4 | positive regulation of neutrophil degranulation | + | 20.83 | 2.58E-05 |
| 5 | cellular response to bacterial lipopeptide | + | 19.44 | 2.15E-04 |
| 6 | cellular response to bacterial lipoprotein | + | 19.44 | 2.15E-04 |
| 7 | response to bacterial lipopeptide | + | 19.44 | 2.15E-04 |
| 8 | cellular response to lipoteichoic acid | + | 17.5 | 7.84E-06 |
| 9 | response to lipoteichoic acid | + | 17.5 | 7.84E-06 |
| 10 | natural killer cell degranulation | + | 17.5 | 1.79E-03 |
| 11 | pentose-phosphate shunt, oxidative branch | + | 17.5 | 1.79E-03 |
| 12 | late nucleophagy | + | 16.66 | 3.28E-04 |
| 13 | pentose-phosphate shunt | + | 14.58 | 9.00E-05 |
| 14 | plasma membrane raft organization | + | 14.58 | 4.80E-04 |
| 15 | response to bacterial lipoprotein | + | 14.58 | 4.80E-04 |

**Supplementary Table 3.** Most relevant enriched gene ontology (GO) terms from genes that are upregulated in nN3 respective to nN2 neutrophils. Terms are organized by fold enrichment, from highest to lowest. Only the 15 biological process GO terms with the highest positive fold enrichment are shown. Raw P value is shown for each term.

| # | GO biological process (upragulated in N3) | +/- | Fold Enrichment | raw P value |
|---|---|---|---|---|
| 1 | nucleolus organization | + | 36.16 | 1.66E-04 |
| 2 | positive regulation of Wnt signaling pathway, planar cell polarity pathway | + | 36.16 | 1.66E-04 |
| 3 | antigen processing and presentation of exogenous peptide antigen via MHC class II | + | 30.14 | 1.94E-06 |
| 4 | synapse pruning | + | 28.93 | 2.83E-04 |
| 5 | antigen processing and presentation of peptide antigen via MHC class II | + | 26.79 | 3.15E-06 |
| 6 | pinocytosis | + | 26.3 | 3.57E-04 |
| 7 | antigen processing and presentation of peptide or polysaccharide antigen via MHC class II | + | 25.38 | 3.95E-06 |
| 8 | antigen processing and presentation of exogenous peptide antigen | + | 17.86 | 1.75E-05 |
| 9 | antigen processing and presentation of exogenous antigen | + | 14.18 | 4.72E-05 |
| 10 | lymphocyte homeostasis | + | 8.77 | 9.72E-05 |
| 11 | antigen processing and presentation of peptide antigen | + | 8.39 | 1.22E-04 |
| 12 | ribosomal small subunit biogenesis | + | 8.15 | 1.41E-04 |
| 13 | leukocyte homeostasis | + | 7.34 | 7.31E-05 |
| 14 | antigen processing and presentation | + | 6.37 | 1.68E-04 |
| 15 | positive regulation of lipid localization | + | 5.87 | 2.70E-04 |

#### SUPPLEMENTARY VIDEOS

**Supplementary Video 1. Neutrophils patrol the extravascular milieu of the nasal mucosa of live uninfected mice.** The nasal mucosa of an uninfected LysM GFP mouse with neutrophils expressing high levels of GFP was visualized using 2-photon microscopy after microsurgery where a burr hole was drilled through the nasal bone to expose the mucosa of the pNR region. Neutrophils appear in green. Vasculature (in white), and cell nuclei (in blue) were visualized after intravascular injection of blood tracer and Hoechst, respectively. Collagen, in blue, was visualized through second harmonics generation (SHG). The dotted line represents the edges of the burr hole. The video is displayed as a time-lapse maximum intensity projection of 3D image stacks generated over 30 minutes, with one stack obtained every 30 seconds for a total of 61 stacks. Scale bar = 30 µm.

**Supplementary Video 2. *Chlamydia muridarum* interacts with neutrophils in the airways.** A LysM GFP mouse was inoculated intranasally with *C. muridarum* dsRed. At 24 hpi, nuclear stain (Hoechst, blue) was injected intravenously and 5 µL of WGA (white) were instilled into the nostrils, to visualize mucus. The mouse was sacrificed 5 minutes after WGA and Hoechst treatment, the NC was dissected and contents of the lumen (i.e. airways) were imaged in situ using 2-photon microscopy. Shown is a 3D image stack approximately 150 µm thick. *C. muridarum* appears in red. Neutrophils appear in green, suspended in mucus (WGA, white), along with other unidentified cells (visible through nuclear staining). To enable visualization of neutrophils and *C. muridarum* alone in this video, WGA and Hoechst signal are eliminated at times 26 sec and 41 sec, respectively.

**Supplementary Video 3.** Neutrophils interact with *Chlamydia muridarum* inclusion bodies in the NM. A LysM GFP mouse was inoculated intranasally with *C. muridarum* dsRed. At 24 hpi, the sample was fixed, stained, and imaged through 2-photon microscopy to generate an 80 µm thick 3D image stack. This video shows a sequence of the optical slices that comprise the stack, through Z. Displayed are neutrophils (green) and *C. muridarum* inclusion bodies (red) that are contained inside epithelial cells. The epithelium (CD326) is shown in white and nuclear stain (DAPI) is shown in blue. To facilitate the visualization of neutrophil-inclusion interactions, the same sequence is shown at time 14 sec without CD326 signal.
